# A New Clustering and Nomenclature for Beta Turns Derived from High-Resolution Protein Structures

**DOI:** 10.1101/390211

**Authors:** Maxim Shapovalov, Slobodan Vucetic, Roland L. Dunbrack

## Abstract

Protein loops connect regular secondary structures and contain 4-residue beta turns which represent 63% of the residues in loops. The commonly used classification of beta turns (Type I, I’, II, II’, VIa1, VIa2, VIb, and VIII) was developed in the 1970s and 1980s from analysis of a small number of proteins of average resolution, and represents only two thirds of beta turns observed in proteins (with a generic class Type IV representing the rest). We present a new clustering of beta turn conformations from a set of 13,030 turns from 1078 ultra-high resolution protein structures (≤1.2 Å). Our clustering is derived from applying the DBSCAN and *k*-medoids algorithms to this data set with a metric commonly used in directional statistics applied to the set of dihedral angles from the second and third residues of each turn. We define 18 turn types compared to the 8 classical turn types in common use. We propose a new 2-letter nomenclature for all 18 beta-turn types using Ramachandran region names for the two central residues (e.g., ‘A’ and ‘D’ for alpha regions on the left side of the Ramachandran map and ‘a’ and ‘d’ for equivalent regions on the right-hand side; classical Type I turns are ‘AD’ turns and Type I’ turns are ‘ad’). We identify 11 new types of beta turn, 5 of which are sub-types of classical beta turn types. Up-to-date statistics, probability densities of conformations, and sequence profiles of beta turns in loops were collected and analyzed. A library of turn types, *BetaTurnLib18*, and cross-platform software, *BetaTurnTool18*, which identifies turns in an input protein structure, are freely available and redistributable from dunbrack.fccc.edu/betaturn and github.com/sh-maxim/BetaTurn18. Given the ubiquitous nature of beta turns, this comprehensive study updates understanding of beta turns and should also provide useful tools for protein structure determination, refinement, and prediction programs.

## Introduction

Ordered protein structures consist of elements of regular secondary structure, such as alpha helices and beta-sheet strands, and irregular elements of structure referred to as loops or coil regions. Loops comprise half of residues in protein structures. Due to the globular nature of folded proteins, the direction of the peptide chain often has to change radically within a few short residues within loops. These changes in direction are often accomplished by turns, which can occur multiple times in loop regions. Turns consist of segments between 2 and 6 amino acids (delta, gamma, beta, alpha and pi turns respectively), and are identified by the distance between the Ca atoms of the first and last residues and sometimes by the existence of specific hydrogen bonds within the segment [1–6]. These turn fragments are often hydrophilic, since loops are usually on the protein surface [7–9].

Four-residue beta turns are by far the most common turn type [5], constituting 25-30% of all protein residues [10]. The first examination of beta turns was by Venkatachalam in 1968 [1]. He identified a key hydrogen bond formed between the backbone carbonyl oxygen atom of the first residue and the backbone amide hydrogen atom of the fourth residue of a beta-turn. Venkatachalam devised a system of three turn types: I, II, and III, and their mirror images I’, II’, and III’. He predicted that the mirror images would be disfavored due to steric clashes.

In 1973, Lewis et al. observed that the backbone 1-4 hydrogen bond is not present in a large number of peptide fragments that might still be considered turns [11]. Instead of the older hydrogen bond criterion, they established the definition of a beta turn requiring a distance between the 1 and 4 alpha-carbon atoms of less than 7 Å, while the central residues 2 and 3 are not part of a helix. This beta-turn definition expansion by Lewis et al. led to the adoption of ten different turn types: I, I’, II, II’, III, III’ from Venkatachalam and new types IV, V, VI, and VII. Type V turns had φ,ψ dihedrals of the 2^nd^ and 3^rd^ residues around (−80°, +80°) and (+80°, −80°) respectively. Type VI turns contained a *cis* proline at the 3^rd^ residue. Type VII turns had y_2_ of 180° or φ_3_ of 180°. The turns not fitting the established definitions, because two or more of the dihedrals of residues 2 and 3 were not within 40° of the defined values, were placed in a miscellaneous category, type IV by Lewis et al.

Richardson et al. [2] kept the same 7 Å criterion used by Lewis et al. and from analysis of a larger data set, reduced the number of beta turn types to six categories with designated φ,ψ limits of the second and third residues: I, I’, II, II’, VIa, VIb, and the seventh miscellaneous category, Type IV. They divided the type VI turns of Lewis et al. with a b conformation at residue 3 and a cis peptide bond at residue 3 into two types: Type VIa with an a conformation at the cis-proline at residue 3 and Type VIb with a b conformation at the cis-proline. They merged type III turns (with φ,ψ =−60°,−30° for residues 2 and 3) with type I turns (with φ_2_,ψ_2_ = −60°,−30° and φ^3^,ψ^3^ =−90°,0° for residue 3) because they occupy similar regions of the Ramachandran map at positions 2 and 3, and many type III turns that could be identified at that time were part of 3_10_ helices. The similarity of 3_10_ helices of length 3 to two consecutive Type I turns has been analyzed by Pal et al. [12]. They found that the first two residues of these short 3_10_ helices have φ,ψ values close to −60°,−30°, while residues 2 and 3 resemble a classical Type I beta turn with φ,ψ=−30°,−60° and −90°,0° respectively.

In 1988 Wilmot and Thornton [3] scanned a dataset of 58 protein structures for turns, and expanded Richardson’s categories by adding type VIII turns, consisting of α and β conformations at the 2^nd^ and 3^rd^ residues respectively [3]. To match each turn to its type, for each residue one of two backbone dihedrals had to be within 30° while the other one was allowed to be within 45° of the canonical values for that turn category. In 1990, the same researchers assigned and named all turns by the Ramachandran plot regions (α_R_, β_E_, β_P_, α_L_, γ_L_, and ε) of the two central residues [4]. They found 16 types that were observed at least once in the total of 910 turns of 58 protein structures. However, eight types were observed less than 10 times (1%) and three types only once. The turn type definitions of Richardson et al. [2] and Wilmot et al. [3] have also guided the design of many turn prediction algorithms [13–19]. For example, in Kountouris et al., the prediction includes types I, II, IV, VIII, and “non-specific” [15].

More recently, de Brevern performed clustering of the miscellaneous beta-turn type IV representing one third of all beta turns [20], those not fitting the criteria for types I, I’, II, II’, VIa1, VIa2, VIb, and VIII [2]. This additional clustering resulted in placing about half of the type IV turns in four groups of turns adjacent to the existing Type I and Type II turn boundaries. However, they do not appear to represent peaks in the density of beta turn conformations, but rather occur at the periphery of Type I and Type II turns after the residues in a 30° × 30° box around the Type I and Type II definitions are removed.

The residue preferences for the most common turn types have been described in previous reports [2, 17, 21, 22]. Hutchinson and Thornton in 1994 analyzed the residue preferences in 2,233 examples of the common turn types in a set of 205 protein chains [17]. They noted the preference for proline at positions in turn types that require f of approximately – 60°, such as position 2 of Type I, Type II, and Type VIII turns, and the preference for Gly, Asn, Asp and Ser when f>0° (position 3 of Type II turns, position 2 of Type II’ turns, and position 2 and 3 of Type I’ turns). Hutchinson and Thornton observed the preference for Asp, Asn and Ser at positions that require a residue in the bridge region of the Ramachandran map (φ =−80°, ψ =0°), for instance at position 3 of Type I and Type II’ turns. Some residues are preferred because they make hydrogen bonds to other residues within the turn or even to the residue before or after the turn. Some hydrophobic residues are preferred when the turn is a tight beta turn connecting two beta-sheet strands, particularly for positions 1 and 4 of Type I’ and II’ turns, in some cases due to their role as part of hydrophobic cores of proteins. They also noted that preference for Pro at position 4 of Type VIII turns, which prevents residue 3 from being in the alpha region of the Ramachandran map [23]. Type VIII turns have a beta conformation residue at this position, while a Type I turn contains an alpha conformation at position 3.

Although many analyses of beta turns have been performed, there are a number of questions that remain outstanding. First, what is the nature of the Type IV beta turns, and can the existing nomenclature be extended to reduce their number? Second, what are the true modes of the density in dihedral angle space for each beta turn? For highly skewed populations, the most common conformation (i.e. modes in the density) for each beta turn would be useful in protein structure determination, refinement, and prediction. Third, what are the amino acid distributions of each beta turn type and can they be rationalized? These have not been reported in detail since the 1990s. Fourth, is the Cα_1_-Cα_4_ distance cutoff of 7 Å justified? And finally, what is the frequency of beta turns in loops of various lengths? This information would be useful in the modeling of longer protein loops, which remains a challenging problem [24].

With a much larger number of ultra-high resolution structures now available, we have undertaken a fresh analysis of the nature of beta turns in protein structures. We have classified all of the beta turn conformations in a data set of 1,078 non-redundant protein chains from high-quality crystal structures of 1.2 Å resolution or higher. In keeping with previous analyses [2, 11], we define beta turns as four-residue segments with the Cα atoms of residues 1 and 4 having a distance of ≤ 7.0 Å and secondary structure of the 2^nd^ and 3^rd^ residues different from sheet (E), helix (H), π-helix (I) and 3_10_-helix (G) according to DSSP [10]. Our clustering is performed with a distance measure based on the seven dihedral angles that connect the Cα atoms of residues 1 and 4: ω_2_, φ_2_, ψ_2_, ω_3_, φ_3_, ψ_3_ and ω_4_, similar to what we used for the clustering of antibody CDRs [25].

To identify clusters that truly represent peaks in the density of data points in the space spanned by these dihedral angles, we have utilized a density-based clustering algorithm, DBSCAN [26]. After optimizing the clustering parameters of DBSCAN, we obtained a set of 11 clusters from the loop residues in our data set of 1,078 proteins. From kernel density estimates of the Ramachandran maps of residues 2 and 3, we identified multiple peaks in the density consistent with the existence of multiple clusters in some of the DBSCAN clusters. We were able to subdivide several of these turn types with *k*-medoids based on the number of peaks in the density, resulting in a set of 18 distinct turn types. We apply a refined division of the Ramachandran map into distinct populated regions developed by Hollingsworth and Karplus [27] to derive a simple nomenclature for our 18 turn types. For our turn types we provide modal conformations and residue preferences at the first, second, third and fourth positions. We analyze the distribution of the Cα_1_-Cα_4_ distance in our beta turn types, and find that some turn types have median Cα_1_-Cα_4_ distances slightly larger than 7 Å, if the 7 Å cutoff is relaxed. We also developed a cross-platform Python script for determining the beta turn types in an input PDB structure.

## Results

### Clustering of a high-resolution data set of beta turns

With the PISCES server [28, 29], we compiled a set of 1,074 protein chains with a resolution equal to or better than 1.2 Å and pairwise sequence identity less than 50% (**S1 Data Set**), comprising protein sequences with 232,197 residues (Table 1). For clustering purposes a total of 16% of all residues were excluded due to missing backbone, Cβ, or γ heavy atom coordinates, chain breaks, presence of alternative conformations, or missing secondary structure information. The resulting data set is named *RefinedSet* with 195,322 residues suitable for turn type analysis. If we define beta turns as four consecutive residues where the central residues 2 and 3 are not in regular secondary structure (alpha helix, 3_10_ helix, π helix or beta strands) and the Cα_1_-Cα_4_ distance is less than or equal to 7 Å, we can locate 13,030 beta turns in this data set built on 43,162 residues. This is less than 13,030 × 4 or 52,110 residues because 49% of beta turns in *RefinedSet* share residues with one or more other beta turns.

**Table 1.** Data sets. Loop, beta-turn, and residue statistics in *CompleteSet* and *RefinedSet*. Sequences contain residues with or without coordinates. *RefinedSet* is a subset of *CompleteSet*, which contains all residues with coordinates. *RefinedSet* excludes residues with multiple conformations (e.g., those with A and B conformations) and those residues with missing backbone, Cβ, or γ heavy atoms. An isolated β turn is a β turn not sharing any of its residues with residues from other β turns.

|  | Count | No of residues | Residue perc. |
| --- | --- | --- | --- |
| Sequences | 1,074 | 232,197 | - |
| *CompleteSet* |  |  |  |
| Residues | - | 224,250 | 100.0 |
| Loops | 17,176 | 102,357 | 45.5 |
| $\beta$ turns | 21,454 | 64,632 | 28.7 |
| Isolated $\beta$ turns | 7,708 | 30,832 | 13.7 |
| *RefinedSet* |  |  |  |
| Residues | - | 195,322 | 87.1 |
| $\beta$ turns | 13,030 | 43,162 | 19.2 |
| Isolated $\beta$ turns | 6,649 | 26,596 | 11.8 |

Beta-turn conformations form overlapping clusters of varying density. Application of a single clustering method such as *k*-means, *k*-medoids, or DBSCAN failed to identify all known turn types and created merged clusters with multiple modes in the density. It would either (1) detect most known turn types but fail to separate turns of Type I and Type VIII or (2) separate these two types as two clusters but in addition produce too many small clusters and noise points. Ultimately, a procedure consisting of applying DBSCAN with optimized parameters followed by *k*-medoids to divide the initial DBSCAN clusters that clearly contained multiple peaks in the dihedral-angle space density proved fruitful.

A single application of DBSCAN on all available 13,030 turns produced 11 clusters with optimized clustering parameters, eps and minPts (Methods). Kernel density estimates (KDEs) of residues 2 and 3 indicated that four of these clusters could productively be subdivided into more than one cluster (Fig 1). The most populous DBSCAN cluster contains 8,455 turns of the classical Type I and VIII turns but in fact exhibits five peaks in the KDE of residue 3 (Fig 1A). Repeated application of *k*-medoids to subdivide this cluster, produced 5 clusters that correspond to well-known features in the Ramachandran map of most residue types, including the alpha–helical region, the gamma-prime or inverse gamma turn region (φ,ψ = −84°,68°) [30], the prePro zeta conformation (φ,ψ = −137°,76°) [31], and two conformations in the beta region, roughly equivalent to parallel (φ,ψ = −120°,125°), and anti-parallel beta sheet (φ,ψ = −134°,157°), regions. Each of these 5 sub-clusters exhibited different amino acid distributions indicating that they are indeed distinct turn types.

**Fig 1.**
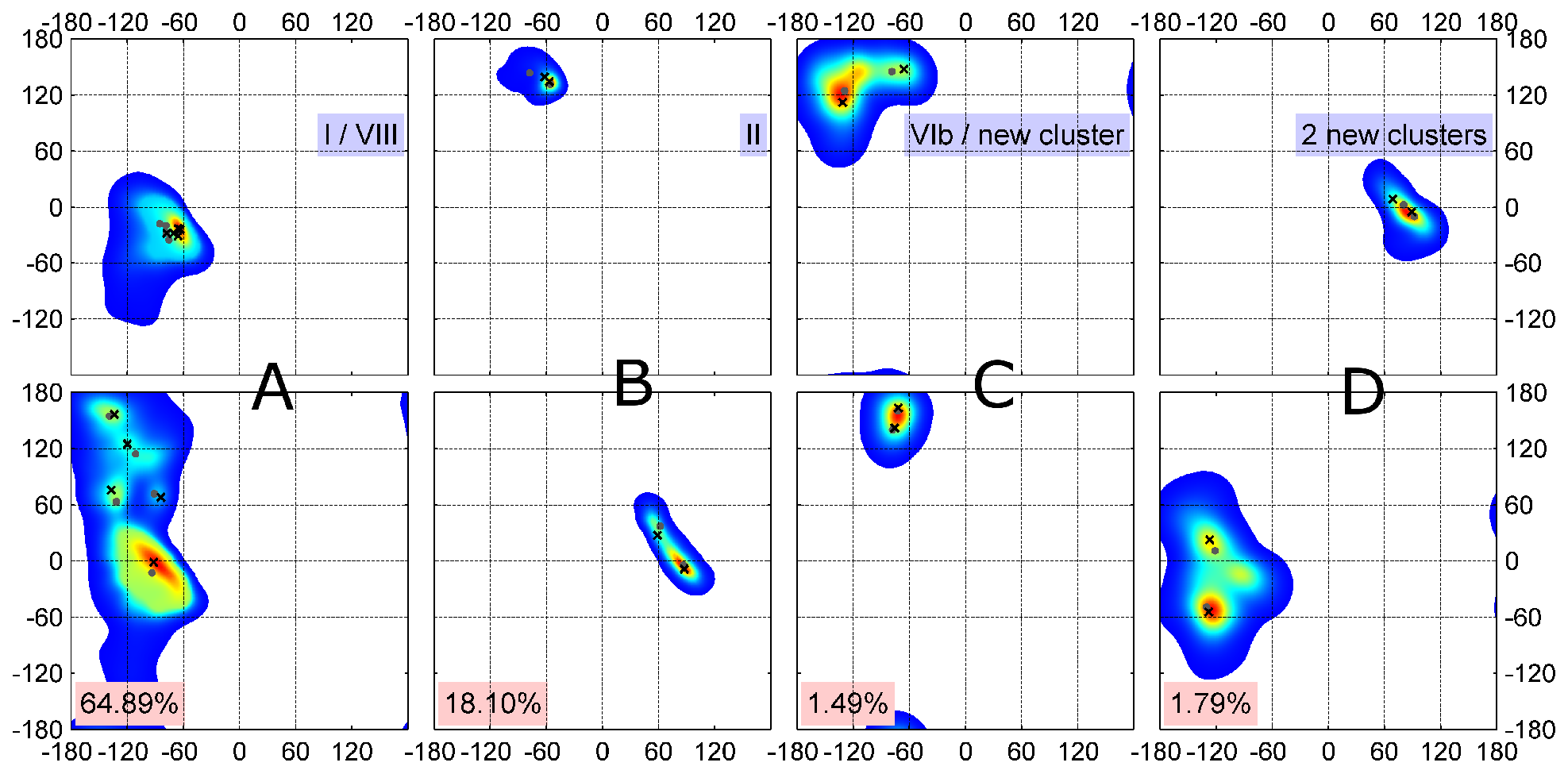
Two-dimensional (φ, ψ) kernel density estimates (KDE) of turn residue 2 (top row) and 3 (bottom row) for four multi-modal clusters returned by DBSCAN (of 11 total). These 4 clusters have multiple peaks in their densities and were further individually divided into subclusters with one or more rounds of *k*-medoids. Their final medoids and modes are marked with gray dots and black crosses respectively. The populations of the four clusters are shown in pink. (A) Cluster for classical Type I and VIII turns exhibit 5 peaks. Type I has the strongest peak and account for 76% of the data points in the cluster. (B) Classical Type II turns can be subdivided into two new clusters; (C) Classical VIb turns with a cis peptide bond at residue 3 can be divided into two types. (D) A DBSCAN cluster with residue 2 in the left-handed helical region and residue 3 adjacent to the right-handed helical region. The data have 3 obvious density peaks but the smallest peak (φ_3_, ψ_3_ ~ −90°,−20°) did not have a distinct amino acid profile and had a very small number of points. We divide this cluster into two beta turn types.

The classical Type II turn cluster from the DBSCAN run contained two peaks, which could easily be subdivided by *k*-medoids at opposite ends of the left-handed alpha helical region in the Ramachandran map for residue 3 (Fig 1B). The upper peak of these conformations preferred glycine at residue 3, while the other was dominated by non-glycine residues at position 3 (see Fig 2 for comparison of amino acid profiles in all clusters). The cluster from DBSCAN in Fig 1C, consisting of some of the structures in *RefinedSet* with a cis peptide bond at residue 3, exhibited two peaks in residue 2 that produced an equivalent to the previously defined Type VIb turn and a new turn type. Finally, DBSCAN produced a new turn type cluster with left-handed helical residues at position 2 and right-handed helical residues at position 3. This cluster contained two dominant peaks in the density for residue 3 (Fig 1D). We subdivided it into 2 separate clusters with *k*-medoids. *k*-medoids is a similar algorithm to k-means but instead selects actual data points as the cluster exemplars instead of a vector of average values in each dimension. In principle, one of these clusters could be further subdivided but we did not find distinct amino acid profiles, and the smaller peak contained a small number of points.

**Fig 2.**
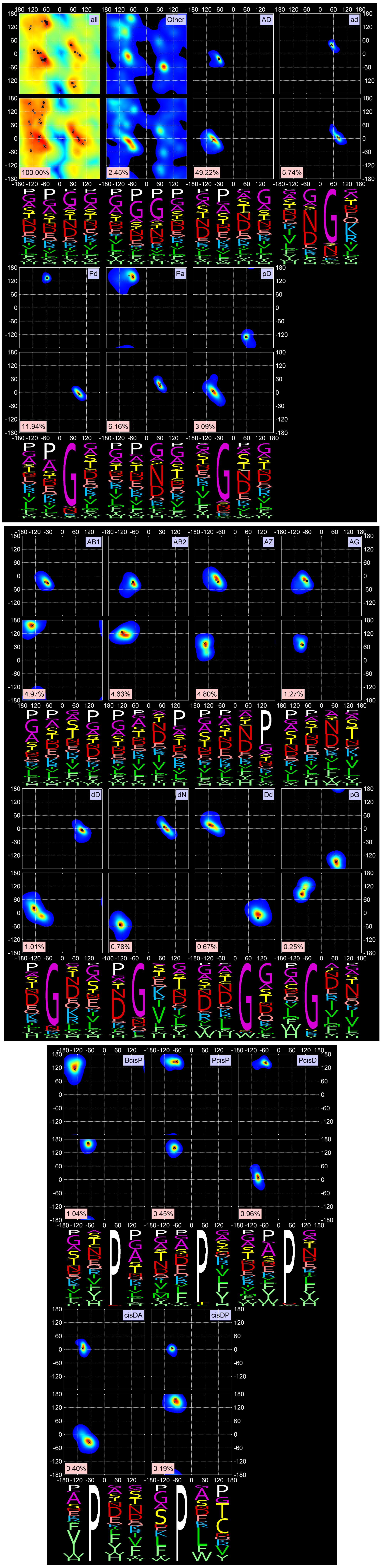
Two-dimensional KDEs for (φ_2_, ψ_2_) and (φ_3_, ψ_3_) and amino-acid sequence profiles for 18 turn types, all turns, and *Other* turns. The mode and medoid of each cluster are shown with a black cross and gray dot respectively. The four-letter profiles are shown for the four amino-acid residues composing a beta turn. First row: All data points, *Other* data points (unclassified), AD (classical Type I) and ad (Type I’) turns. “All” data points KDE is shown on a log scale. Second row: Pd and Pa turns (Type II) and pD turns (Type II’). Third row: AB1, AB2, AZ, and AG turns (all formerly Type VIII). Fourth row: new turn types dD and dN and their approximate inverse, Dd, and new turn type pG. Fifth row: cis3 turn types, BcisP (Type VIb), PcisP (new), and PcisD (Type VIa1). Sixth row: new cis2 turns, cisDA and cisDP.

Our clustering procedure results in a set of 18 turn types (Table 2 and Fig 2) among the 13,030 beta turns detected in *RefinedSet*. We define the noise points identified by DBSCAN as a group called *“Other*,” which consists of 319 turns or 2.4%. Table 2 contains the populations among our 13,030 beta turns and the modal values of φ and ψ for residues 2 and 3. Among our 18 turn types and the *Other* group: 6 types are new; 8 types are created by splitting existing Type II turns (2 types), Type VIII turns (4 types), and Type VIb turns (2 types); and the remaining 4 types are updated versions of the remaining classical turn types (I, I’, II’, VIa1). We did not find a cluster of the classical VIa2 type with a cis peptide bond at residue 3 and φ_2_,ψ_2_ = [−120°,120°] and φ_3_,ψ_3_ = [−60°,0°]. Instead, a small number of points near these values occur in the *Other* group, but they are too spread out in dihedral angle space and not numerous enough to produce a cluster. By contrast, we do find two types within a cis peptide bond and a β-like conformation at residue 3. One of these is close to the classical VIb turn; the other is new.

**Table 2.** A set of 18 beta turn types derived from clustering of the *RefinedSet* of beta turns. The new nomenclature (column 4) based on the Ramachandran map positions (Figure 3) of residues 2 and 3 is provided next to the classical nomenclature. “cis” precedes conformations that contain cis peptide bonds for ω_2_ or ω_3_. “New” turns are those not similar to any types in the classical nomenclature. Some new turns are subclusters of the previous turn types; the one closest to the classical turn type is labeled with that type; the others are labeled “new” with the previous parent turn type given. The modal values of the dihedrals φ and ψ and the cis/trans configuration of residues 2 and 3 are shown. Complete information for 18 beta turns is provided in S1 Table and S1 Text. The *BetaTurnLib18* files and beta-turn assignment Python tool, *BetaTurnTool18* are at http://dunbrack.fccc.edu/betaturn or http://github.com/sh-maxim/BetaTurn18.

| No.<br>by<br>size | Size | Perc. | New<br>name | Previous name | Median<br>Ca<sub>1</sub>-Ca<sub>4</sub><br>distance | Residue 2<br>modal value |  |  | Residue 3<br>modal value |  |  |
| --- | --- | --- | --- | --- | --- | --- | --- | --- | --- | --- | --- |
| | | | | | | $\omega$ | $\phi$ | $\psi$ | $\omega$ | $\phi$ | $\psi$ |
| 1 | 6,413 | 49.22 | AD | I | 5.48 | *trans* | -62 | -23 | *trans* | -96 | -2 |
| 2 | 1,556 | 11.94 | Pd | II | 5.61 | *trans* | -55 | 133 | *trans* | 91 | -6 |
| 3 | 802 | 6.16 | Pa | new (prev. II) | 6.00 | *trans* | -60 | 135 | *trans* | 59 | 28 |
| 4 | 748 | 5.74 | ad | I' | 5.44 | *trans* | 47 | 45 | *trans* | 83 | 1 |
| 5 | 648 | 4.97 | AB1 | new (prev. VIII) | 6.72 | *trans* | -67 | -31 | *trans* | -136 | 162 |
| 6 | 625 | 4.80 | AZ | new (prev. VIII) | 5.80 | *trans* | -74 | -28 | *trans* | -140 | 75 |
| 7 | 603 | 4.63 | AB2 | VIII | 6.42 | *trans* | -69 | -30 | *trans* | -120 | 128 |
| 8 | 402 | 3.09 | pD | II' | 5.44 | *trans* | 57 | -130 | *trans* | -95 | 11 |
| 9 | 166 | 1.27 | AG | new (prev. VIII) | 6.37 | *trans* | -66 | -19 | *trans* | -82 | 63 |
| 10 | 135 | 1.04 | BcisP | VIb | 5.70 | *trans* | -138 | 119 | *cis* | -66 | 164 |
| 11 | 131 | 1.01 | dD | new | 6.63 | *trans* | 94 | -1 | *trans* | -128 | 15 |
| 12 | 125 | 0.96 | PcisD | VIa1 | 5.74 | *trans* | -60 | 144 | *cis* | -93 | 8 |
| 13 | 102 | 0.78 | dN | new | 6.42 | *trans* | 69 | 9 | *trans* | -132 | -63 |
| 14 | 87 | 0.67 | Dd | new | 6.81 | *trans* | -115 | 16 | *trans* | 101 | -12 |
| 15 | 59 | 0.45 | PcisP | new (prev VIb) | 6.33 | *trans* | -66 | 148 | *cis* | -76 | 142 |
| 16 | 52 | 0.40 | cisDA | new | 5.36 | *cis* | -94 | 8 | *trans* | -61 | -38 |
| 17 | 32 | 0.25 | pG | new | 6.58 | *trans* | 74 | -162 | *trans* | -79 | 77 |
| 18 | 25 | 0.19 | cisDP | new | 6.36 | *cis* | -86 | 4 | *trans* | -71 | 158 |
| other | 319 | 2.45 | Other | IV | 6.42 |  |  |  |  |  |  |
| **Total** | **13,030** | **100.00** |  |  |  |  |  |  |  |  |  |

Rather than the Roman numeral system in use since 1973 with abandoned numerals (e.g., III, V, and VII) and a set of intricate prime symbols and alphanumerical sub-indices (e.g., I’, II’, VIa1), we propose a new nomenclature using the Latin alphabet with type names assigned according to the one-letter region codes of Ramachandran map for the 2^nd^ and 3^rd^ residues of each beta turn type (Fig 3). The figure shows our designations and the positions of the medoids of residues 2 and 3 (orange dots).

**Fig 3.**
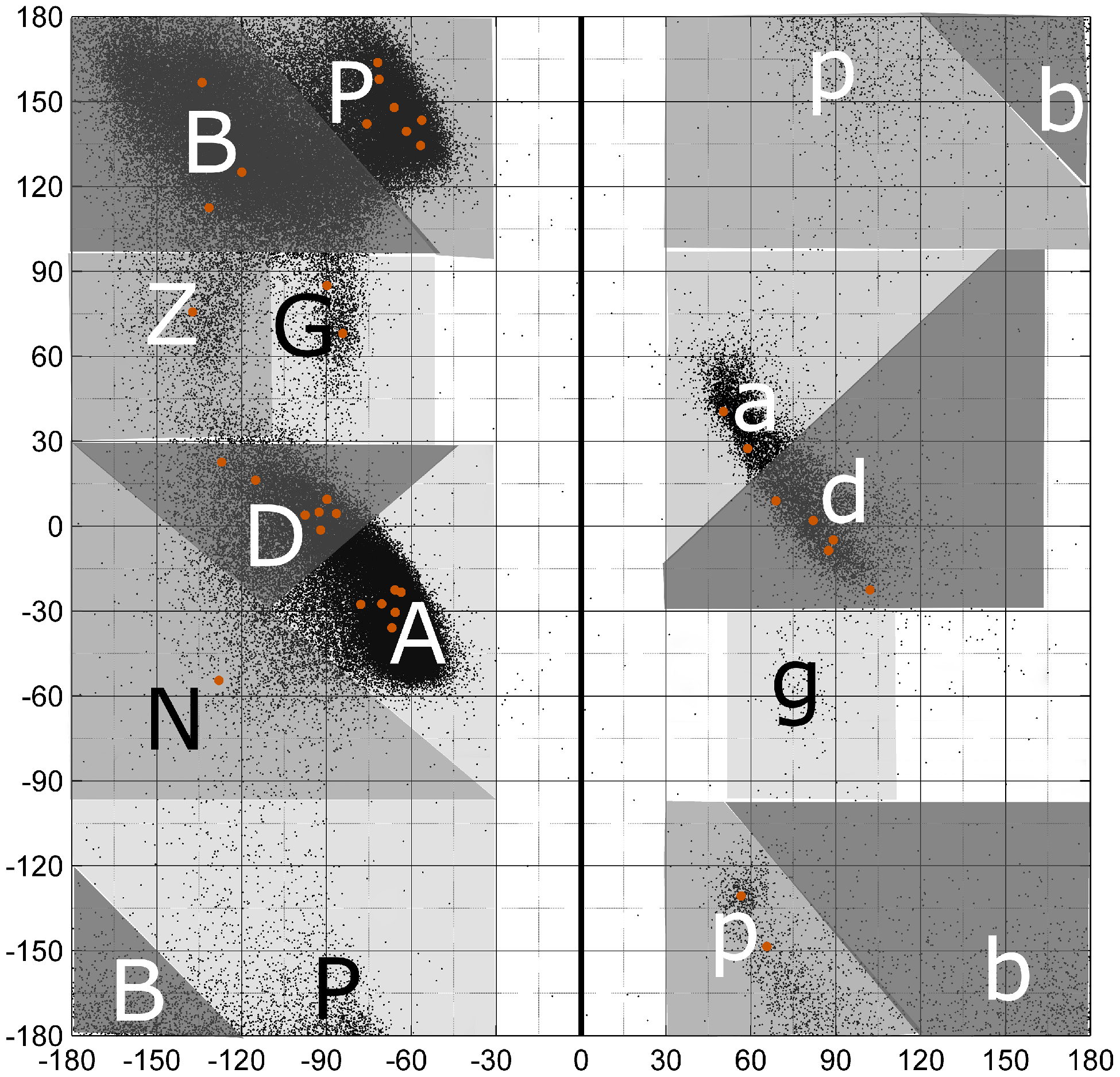
Single-letter region codes for the Ramachandran map. Greek letter designations were converted to Roman letter counterparts. The left side of the map with negative φ is encoded in upper-case letters, while the right side of the map with positive φ is in lower-case letters. Regions related by symmetry through the origin are denoted with the same letter (e.g., A and a; P and p). The original gamma and gamma prime regions are labeled g and G respectively. The new label N is used for the region below the A-D axis. All cluster medoids of residues 2 and 3 are marked with orange dots. The black dots represent data from all amino acids in the *RefinedSet*, including regular secondary structures and loops.

Our scheme is based on a detailed analysis of the distribution of residues from very high-resolution structures by Hollingsworth and Karplus [27]. Most of the one-letter Ramachandran codes are familiar to structural biologists. For convenience both in computer code and for people, we converted their Greek designations to the Latin alphabet, for example α to A, β to B, γ to G and so on. We reserve upper-case letters for the left side of Ramachandran map with negative φ and lower-case letters for positive φ. The upper and lower case letters have a point symmetry relative to the map center (0, 0). This is similar to nomenclatures developed by Dasgupta et al. [32] and Hollingsworth et al. [33] A non-standard designation in this scheme is to denote the gamma turn region at φ = −84°, ψ = 68° as “G” in upper case. Originally, this region was designated gamma-prime (G’ or g) and the corresponding region with φ > 0° was designated as gamma (G). We also defined a new region, N, to the left and below the A-D density axis, which occurs in the dN cluster. This “plateau” region was determined to be sparsely populated but valid by Lovell et al. [34]. In only one case (AB1 and AB2) do we need a third character to distinguish the two turn types. AB1 is the upper one the Ramachandran map and AB2 is the lower one (Fig 1A and Fig 2), which is an easy way to remember them. In Table 2, we also provide a correspondence between the proposed and classical turn type nomenclatures.

Fig 2 contains the amino acid distributions at all four positions of each beta turn type, including the *Other* group. As expected, glycine and proline play important roles in many turn types. Proline is required for the two cis2 turn types, cisDA and cisDP, and the three cis3 turn types, BcisP, PcisP, and PcisD. It is also important in position 4 of the AZ turn where it is responsible for the zeta conformation at position 3, which occurs for residues immediately before proline [27]. Proline also prevents the alpha conformation at the residue before it [23], and is therefore significant in the AB1 and AB2 turns at position 4 and completely absent from AD turns at positions 3 and 4.

Glycine allows amino acids to occupy the “d” and “p” regions of the Ramachandran map, as exhibited by the ad, Pd, dD, dN, and pG type turns. As noted above, the former Type II turns have been divided into Pa and Pd; glycine is dominant at position 3 in the Pd cluster but not in the Pa cluster, which is instead dominated by Asn and Asp at position 3. Asp and Asn are also dominant at position 2 of the ad turn (former Type I’) and position 3 of the AG and AZ turns.

There are sometimes subtle but statistically significant differences in the sequence profiles of very similar clusters. For instance, residue 3 of AB1 and AB2 has slightly different positions in the Ramachandran map. The upper peak (AB1) prefers Gly, Ala, Ser, and Thr at position 3, while the lower peak (AB2) prefers Asn and Asp. The dD and dN clusters are different at position 3, with dD preferring Asn and Asp, which are almost absent from position 3 of the dN turns. The latter contain hydrophobic beta-branched residues at position 3 (Val and Ile).

The new turn types, PDB entry, chain, and residue identifiers, dihedral angles of medoids and modes, and secondary structure context are provided in *BetaTurnLib18* turn library: S1 Text (text format) and S1 Table (Excel format).

### Validation and structural properties of the new beta turn types

We validated the new turn types by examining their electron density both visually and quantitatively. To analyze the electron density of atoms in beta turns, we utilized the EDIA program (Electron Density Support for Individual Atoms) [35], which integrates electron density in a sphere around each heavy atom and penalizes both positive and negative density in electron density difference maps. EDIA demonstrates that turns in each cluster contain only a few structures (0-2.4%) with substantial inconsistencies (Fig 4). The *Other* group has a higher rate of substantial inconsistencies with 6.1% of these turns having poor electron density. Nevertheless, the majority of *Other* group turns are valid conformations, but sufficiently spread-out that they do not form a new cluster with at least 25 points (0.2% of *RefinedSet)*. The mean EDIA scores for each cluster are provided in Table 3, where the EDIA score for each turn is defined as the *minimum* value for the backbone atoms defining the turn, i.e. atoms from Cα_1_ to Cα_4_, including the carbonyl oxygens.

**Fig 4.**
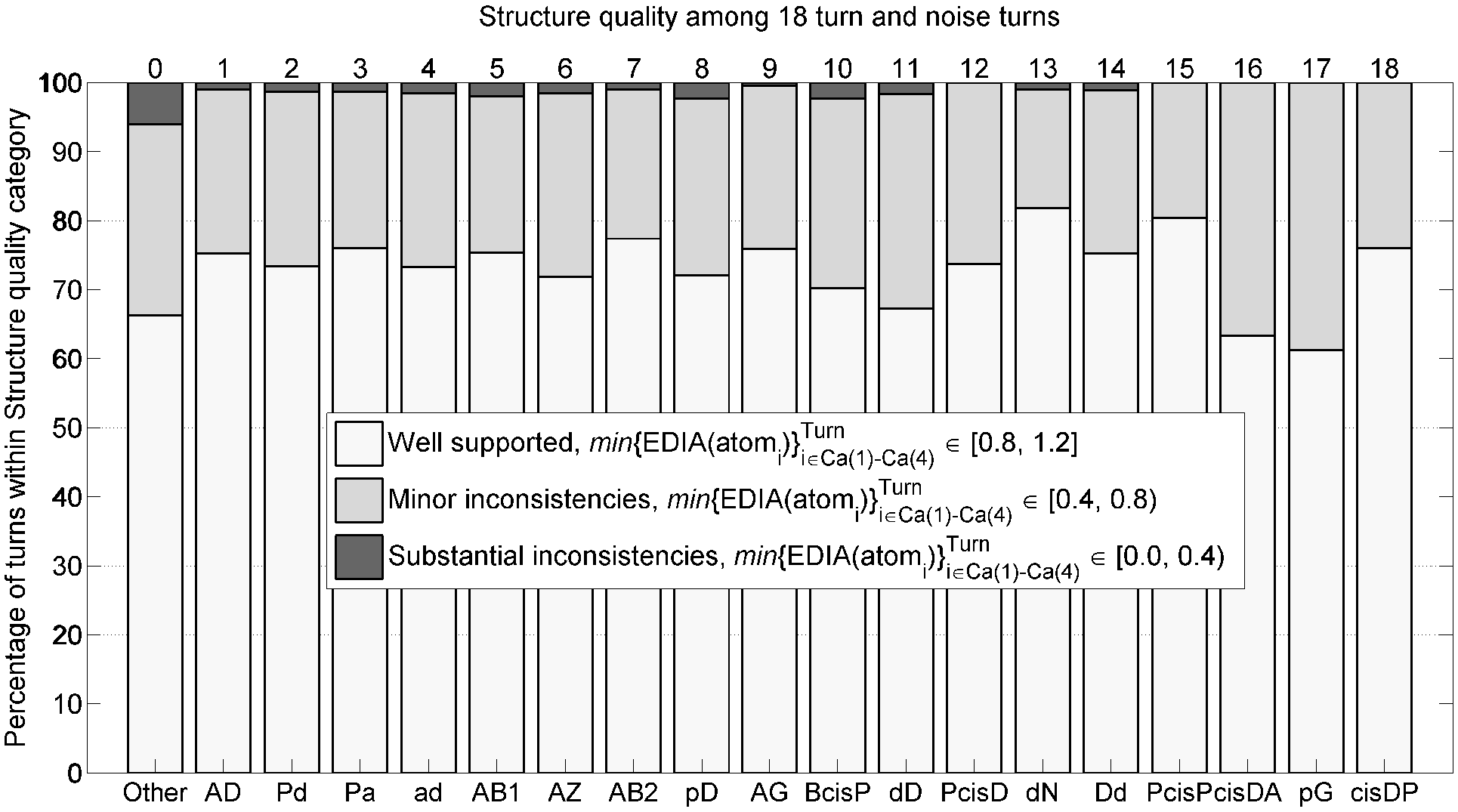
Structure quality of *Other* turns and 18 turn types based on agreement between atom positions and electron-density maps measured by EDIA software with the minimum of *EDIA* values for all 13 backbone atoms (N, Cα, C, O) in a turn connecting Cα1 to Cα4.

**Table 3.** Validation of beta turn clusters. Column 3: Mean EDIA scores for the minimum EDIA score for all backbone atoms of each beta turn in each cluster. Beta turns in *RefinedSet* were reassigned with the *BetaTurnTool18* program with the standard dihedral angle cutoff and the 7 Å Cα_1_-Cα_4_ cutoff. Columns 4-11: Reassignments based on these criteria are provided, consisting of counts of “Original cluster size”, number lost to another regular cluster (1-18), number lost to *Other*, number reassigned from another regular cluster and number reassigned from *Other*. The total number of reassigned turns in each cluster is given in column 9. The TPR (true positive rate) is defined as the percent of the assignments that are present in the reassignments, which is equal to 100*[(Column 4) + (Column 5) + (Column 6)] / (Column 4); PPV (positive predictive value) is defined as the percent of the reassignments that were present in the original assignments, which is equal to 100*[(Column 4) + (Column 5) + (Column 6)]/ (Column 9). It is the percent of turns in the reassigned set that come from the same cluster in the original set. Column 11: Median Cα_1_-Cα_4_ distance for each cluster if the *BetaTurnTool18* program is applied with the standard dihedral angle distance cutoff but no Cα_1_-Cα_4_ distance cutoff. Segments with dihedral angle distance larger than the cutoff are assigned to *Other;* most of which are not turn-like structures.

| No. | New name | EDIA mean | Original Size | Lost to another cluster | Lost to *Other* group | Gained from other clusters | Gained from *Other* group | New Count | TPR | PPV | Median Ca<sub>1</sub>-Ca<sub>4</sub> distance, Å (no cutoff) |
| --- | --- | --- | --- | --- | --- | --- | --- | --- | --- | --- | --- |
| 1 | AD | 0.854 | 6,413 | -149 | -195 | +0 | +0 | = 6069 | 94.6 | 100.0 | 5.51 |
| 2 | Pd | 0.845 | 1,556 | -1 | -0 | +0 | +6 | = 1561 | 99.9 | 99.6 | 5.61 |
| 3 | Pa | 0.848 | 802 | -5 | -30 | +0 | +2 | = 769 | 95.6 | 99.7 | 6.12 |
| 4 | ad | 0.839 | 748 | -0 | -3 | +0 | +3 | = 748 | 99.6 | 99.6 | 5.45 |
| 5 | AB1 | 0.851 | 648 | -1 | -12 | +0 | +0 | = 635 | 98.0 | 100.0 | 7.52 |
| 6 | AZ | 0.845 | 625 | -47 | -1 | +128 | +0 | = 705 | 92.3 | 81.8 | 5.90 |
| 7 | AB2 | 0.855 | 603 | -11 | -7 | +56 | +0 | = 641 | 97.0 | 91.3 | 7.95 |
| 8 | pD | 0.836 | 402 | -9 | -4 | +0 | +5 | = 394 | 96.8 | 98.7 | 5.66 |
| 9 | AG | 0.855 | 166 | -14 | -1 | +38 | +0 | = 189 | 91.0 | 79.9 | 7.24 |
| 10 | BcisP | 0.832 | 135 | -0 | -0 | +0 | +2 | = 137 | 100.0 | 98.5 | 6.07 |
| 11 | dD | 0.821 | 131 | -5 | -2 | +5 | +10 | = 139 | 94.7 | 89.2 | 7.79 |
| 12 | PcisD | 0.858 | 125 | -0 | -0 | +0 | +8 | = 133 | 100.0 | 94.0 | 5.76 |
| 13 | dN | 0.873 | 102 | -4 | -0 | +5 | +1 | = 104 | 96.1 | 94.2 | 7.83 |
| 14 | Dd | 0.856 | 87 | -0 | -0 | +6 | +7 | = 100 | 100.0 | 87.0 | 7.60 |
| 15 | PcisP | 0.861 | 59 | -0 | -0 | +0 | +2 | = 61 | 100.0 | 96.7 | 6.74 |
| 16 | cisDA | 0.827 | 52 | -0 | -0 | +0 | +4 | = 56 | 100.0 | 92.9 | 5.36 |
| 17 | pG | 0.806 | 32 | -0 | -0 | +8 | +4 | = 44 | 100.0 | 72.7 | 8.16 |
| 18 | cisDP | 0.884 | 25 | -0 | -0 | +0 | +11 | = 36 | 100.0 | 69.4 | 6.52 |
| Other | *Other* | 0.802 | 319 | -65 | - | +255 | - | = 509 | 79.6 | 49.9 | 9.03 |
| **Total** |  |  | **13,030** | **-311** | **-255** | **+501** | **+65** | **= 13,030** | **95.7** | **95.7** | **-** |

The electron densities of representative structures near the modes in the seven-dimensional probability distribution are shown for all 18 beta turn types in Fig 5. In addition, the figure shows two structures from the *Other* group that have good electron density, demonstrating that turns in the *Other* group are valid conformations. These were chosen to be far away from any of our cluster modes. Both of these structures have a cis proline at residue 4, but different Ramachandran regions for residues 2 and 3 from each other. There are 21 turns with a cis proline at residue 4 in the *Other* category. They do not cluster well with our seven-dimensional dihedral angle metric.

**Fig 5.**
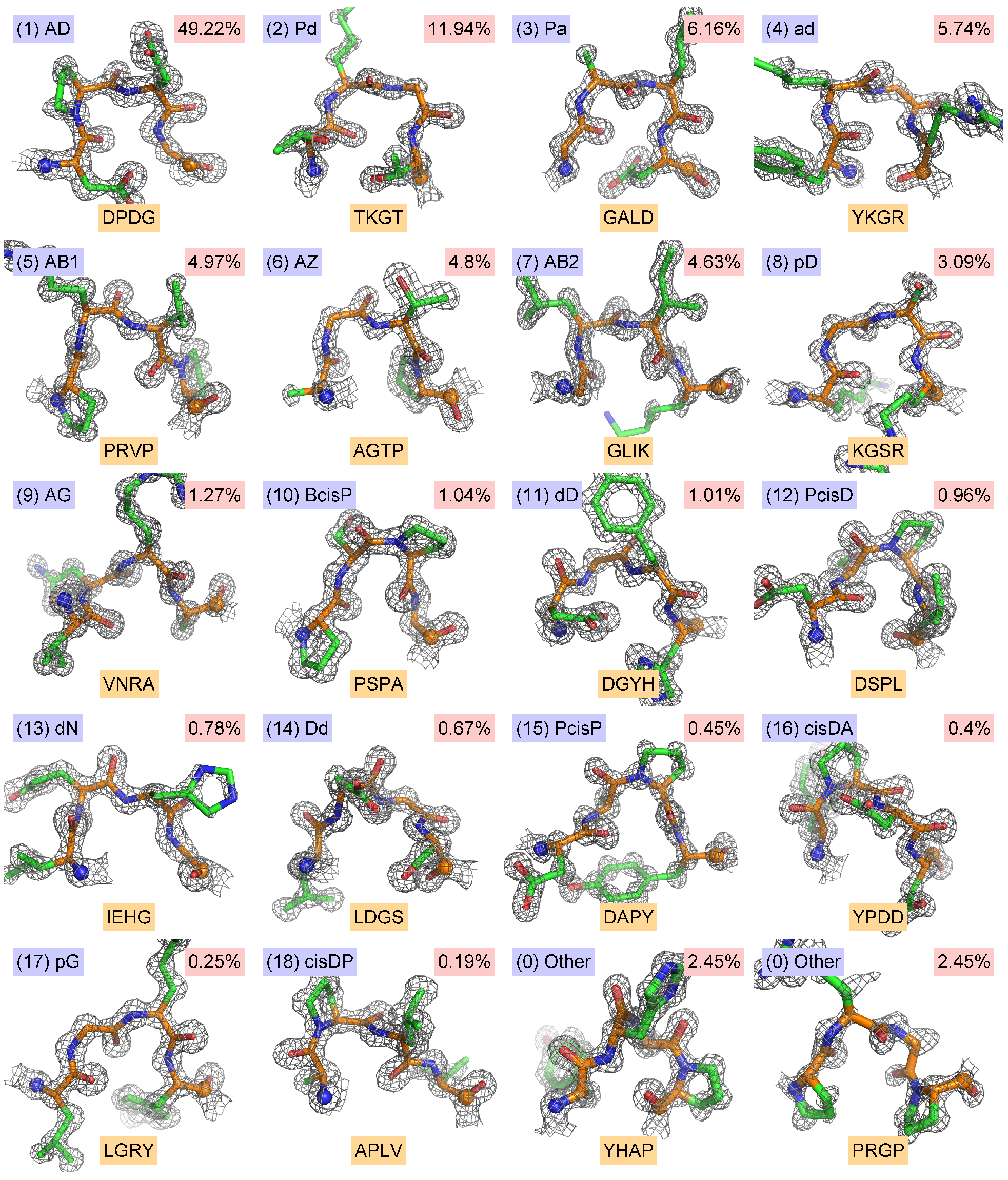
2Fo-Fc electron density distributions for each of the 18 new beta turn types, plus two examples from the *Other* category. The PDB entries and other data for each turn are given in *BetaTurnLib18* (S1 Table and S1 Text). Contours are shown at 2σ. The two *Other* examples (bottom row, 3^rd^ and 4^th^ images) are: (1) PDB entry 2EAB, chain A, residues 759-762, sequence YHAP, conformation APcis4, [φ_2_, ψ_2_; φ_3_, ψ_3_]=[−92°,−39°; −76°,165°]; (2) PDB entry 4RJZ, chain A, residues 366-369, sequence PRGP, conformation Bbcis4, [φ_2_, ψ_2_; φ_3_, ψ_3_]=[−78°,120°; 82°,−127°]. Residue 4 is a cis proline in both cases.

The results of clustering procedures can be validated with a number of measures [36]. Most such measures compare the distances of points within clusters to the distances between points in different clusters in some way. Density-based clustering presents some challenges for standard validation measures, since clusters may be shaped irregularly and have different densities and variances about their medoids compared to distance-based clustering algorithms such as k-means [37]. Nevertheless, we utilized the silhouette score [38] which compares the average distances of points to other points in the same cluster to the average distance of points to the nearest neighboring cluster for each point.

For each point *i*, we calculate the average distance to the points in each cluster (one value for each cluster including the cluster *i* belongs to). If *a*(*i*) is the average distance to points in the same cluster as point *i* and *b(i)* is the distance to the nearest cluster different from that of point *i*, then the silhouette score is defined as *S*(*i*) = (*b*(*i*) – *a*(*i*))/*max*(*a*(*i*),*b*(*i*)). If point *i* is near other points in its own cluster, but very far away from the nearest other cluster, then *S(i)* is close to 1.0. If point *i* is equally close to its own cluster and another cluster, *S(i)* is close to 0, and if point *i* is closer to points in another cluster than its own cluster, *S(i)* is negative. This can occur if point *i* is misplaced or if point *i’s* cluster has a high variance.

A graph of silhouette scores for our 18 clusters and the *Other* group is shown in Fig A **in S1 Supporting Information**. Each of our clusters has an average positive silhouette score ranging from 0.27 to 0.93. Eleven clusters have values of 0.6 or higher. Some of the clusters that arose from sub-clustering of the original DBSCAN clusters with *k*-medoids have relatively low values of the average silhouette score, including Pa (0.27), AB2 (0.29), and AZ (0.32). This is not surprising, since they are very close to other clusters (Pd, AB1/AZ, and AB2/AD/AZ respectively), but as shown in Fig 1, these clusters represent peaks in the density of the Ramachandran maps of residues 2 and/or 3, and they have distinct sequence profiles. The silhouette score has difficulty in scoring sub-clusters that are close together but far from other clusters [36]. We repeated the silhouette analysis by merging AB1 with AB2, AZ with AG, Pd with Pa, and dD with dN to see whether the subclusters are distinct from the rest of our clusters even if they are near related subclusters (Fig B **in S1 Supporting Information**). The AB1-AB2, AZ/AG, Pd/Pa, and dD/dN merged clusters now have higher average silhouette scores of 0.49, 0.48, 0.80, and 0.70 respectively.

The robustness of our clusters may also be assessed by reassigning points in our data set to the cluster with the closest medoid to that point. If we define a distance cutoff for assigning points to a cluster or the *Other* category, and if points are mostly assigned to the correct cluster, then our cluster medoids can be used successfully to assign beta turns in structures not in our data set, including new structures deposited in the PDB. To detect beta turns, we developed a Python script that either assigns one of 18 turn types based on the closest cluster medoid or flags it as *Other* if the distance between the input turn and the closest medoid is above a preselected distance threshold. This maximum distance is equal to the mean plus 3 standard deviations of 12,711 distances between turns of one of 18 types (*Other* excluded) to their cluster medoids, which is equal to 0.2359 in units of our distance metric or 28.1° in units of the average angle equivalent of our metric. We tried different statistics, target thresholds and individual thresholds for each of 18 turn types; a simple single threshold at 3 standard deviations works as well as cluster-specific means and standard deviations in terms of reclassifying beta turns in our set of structures. The results are presented in Table 3.

To understand how the assignment tool performs, we need to account for how the original assignments of turn types of 13,030 beta turns are transformed by reassignment. Most turns are reassigned to the same turn type: (13,030-311-255)/13,030=95.7%. Some turns will be reassigned to one of the other 18 clusters (311); some assigned turns will be reassigned to *Other* (255); some turns will be reassigned from one of the other 18 clusters (256); and some turns will be reassigned from the *Other* category (65). All of these numbers are presented in Table 3 for each cluster and the *Other* category.

We can assess the assignment tool with for each cluster in the way that binary predictors are evaluated. We calculated true positive rate (TPR) as the fraction of the original clustering that is preserved in the reassignments. This is equal to 100.0*(Column_4 + Column_5 + Column_6) / (Column_4). The TPR values range from 91% to 100% for the regular clusters, and 79.6% for the *Other* category. We calculated the positive predictive value (PPV) as the percentage of turns in the reassigned set that came from the original assignments. From the table, this is equal to 100.0*(Column_4 + Column_5 + Column_6) / (Column_9). For 12 of our 18 clusters, this value is well over 90%. It is lower for clusters that are close to other clusters, widely dispersed, or very small clusters, including AZ (81.8%), AG (79.9%), pG (72.7%), and cisDP (69.4%). The value is 49.9% for the *Other* category because of the number of points moved from regular clusters into the *Other* group (255). These points are far from the medoids of non-spherically shaped clusters. They represent only 2% of all the data points.

Finally, as a part of validation of the clusters, since a high threshold of 1.2 Å crystal structure resolution was applied to *RefinedSet* in order to produce more reliable data, we used the tool to determine whether there is a distribution bias in the established turn types as a function of resolution. The frequencies for the most turn types remained the same when proteins at 1.0-1.2 Å and 2.0-2.2 Å (Fig 6) are compared. The relative frequencies of some turn types do change with resolution: Pd (9.7% to 8%), ad (5.0% to 4.0%), AG (1.3% to 2%), and *Other* turns (3.2% to 4.5%)

**Fig 6.**
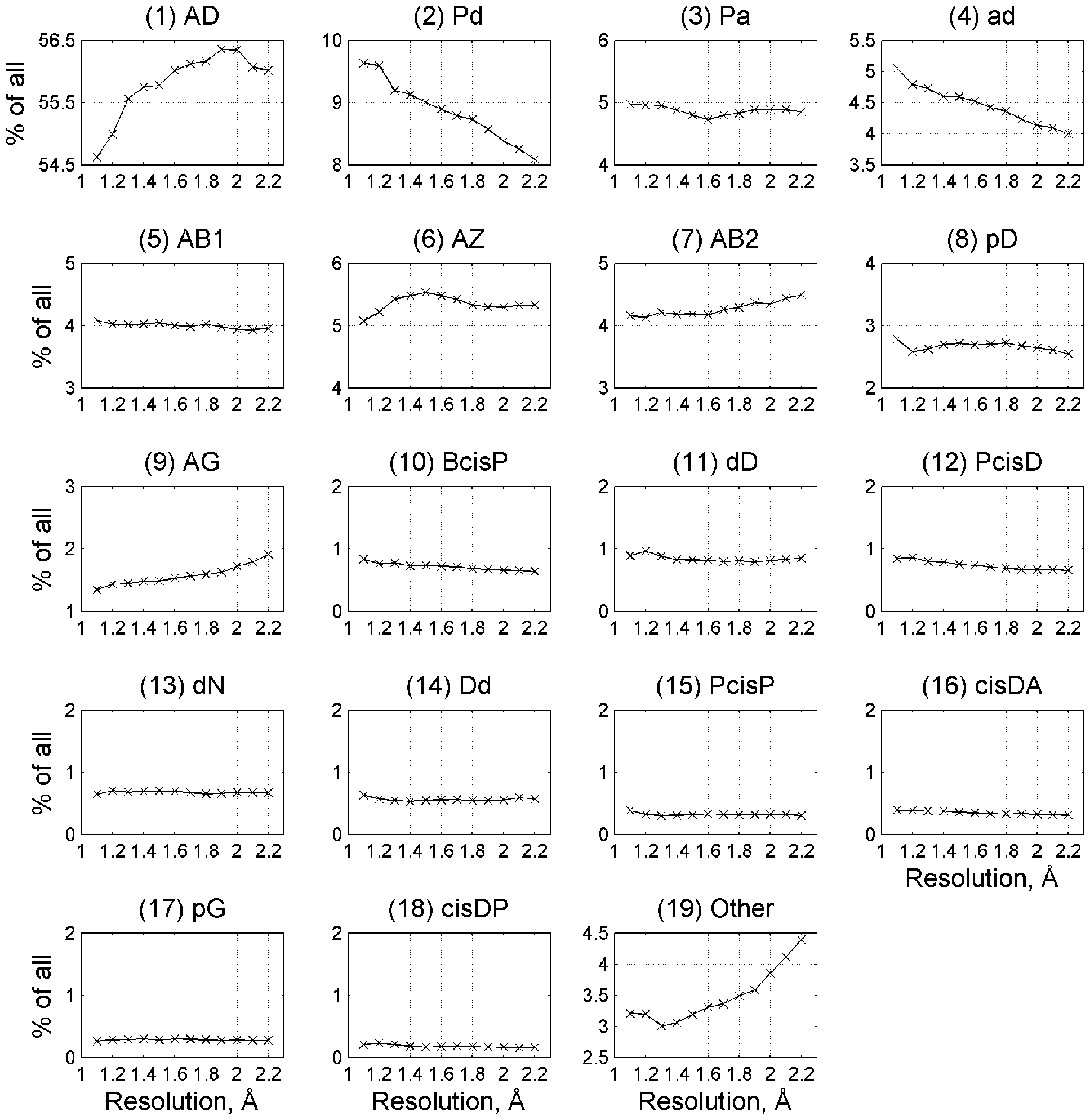
Frequency of beta turn types at different resolutions. Beta turns were identified in structure sets at different resolution ranges (1-1.2 Å, 1.1-1.3 Å, etc.) produced by the PISCES webserver. The y-axis on each plot covers 2 percentage points in total frequency of each beta turn type.

### *Analysis of the* Cα_1_-Cα_4_ *distance in the definition of beta turns*

We used a value of 7.0 Å for the Cα_1_-Cα_4_ distance to define beta turns within our data set. This value has been used in almost all previous studies on beta turns [2, 4, 11, 17, 20, 39]. It is slightly larger than the minimum at about 6.5 Å between two peaks on the distribution of Cα_1_-Cα_4_ distances in all four-residue segments in loops in our data set (Fig 7, bottom right). In Table 2, we provide the median Cα_1_Cα_4_ distance for each of our clusters when beta turns are defined with the classical 7 Å Cα_1_Cα_4_ cutoff. However, we were curious if some beta turns might have a wider distribution of this distance if a larger cutoff or no distance cutoff is used, while maintaining a turn-like conformation. We assigned all four-residue segments in our data set of loops (those residues not in regular secondary structures) to one of our clusters – to the one with the closest medoid, if the distance to the closest medoid was less than or equal to the cutoff distance described above (0.2359 in units of our distance metric). If the distance was greater than this value, we put the segment into the *Other* category. Kernel density estimates of the Cα_1_-Cα_4_ distance when no Cα_1_-Cα_4_ cutoff is applied are shown in Fig 7 for all of our 18 clusters, the *Other* group (four-residue segments not near one of our cluster medoids in dihedral angle space), as well as *All* four-residue segments in the loop data set (the union of all 18 clusters and the *Other* group).

**Fig 7.**
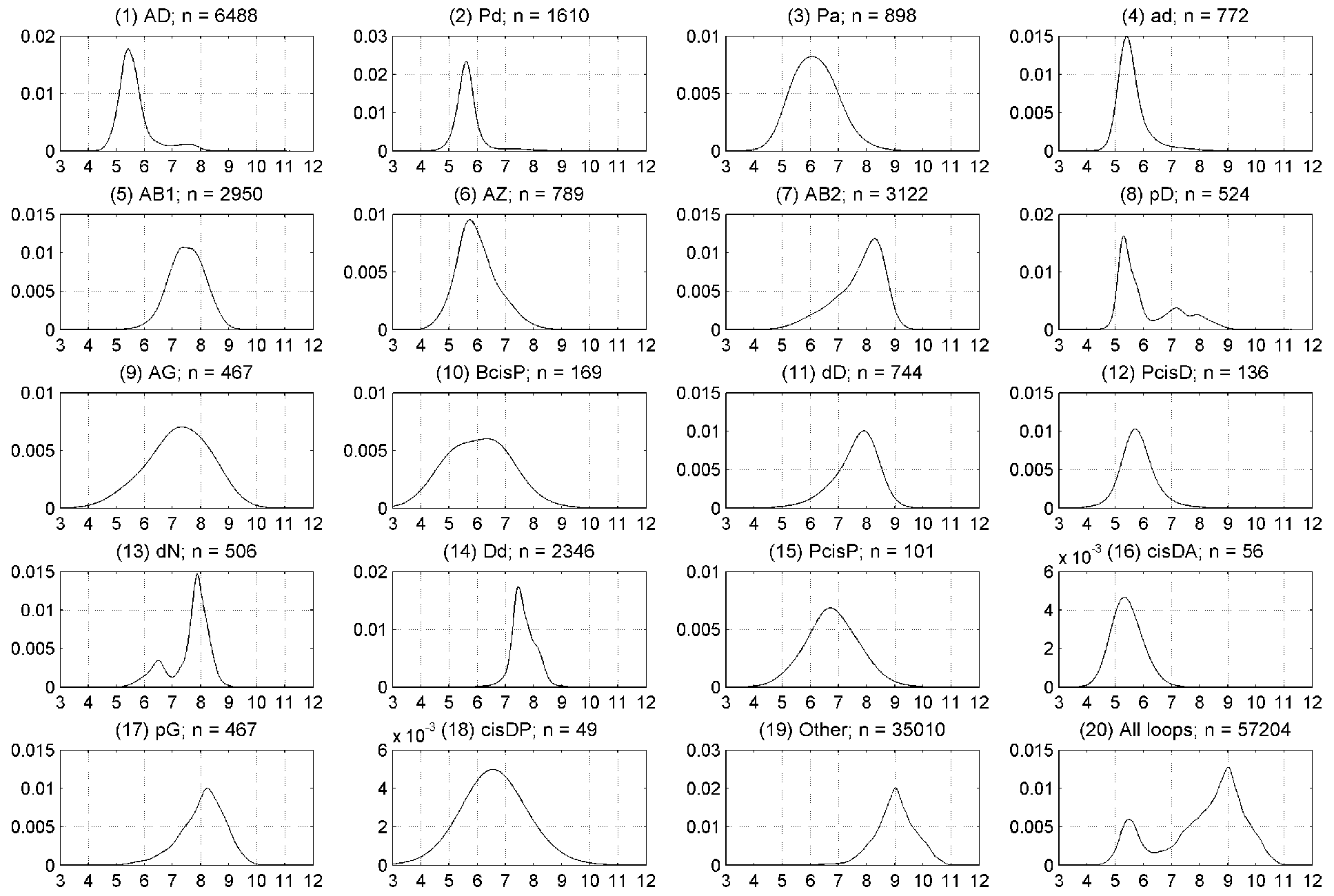
Kernel density estimates of Cα_1_-Cα_4_ distances when no 1-4 distance cutoff is applied. Plots are shown for each of 18 beta turn clusters, the *Other* group, and all four-residue segments in loop regions of the *RefinedSet* (bottom right).

The maximum possible Cα_1_-Cα_4_ distance is about 11.4 Å (3 × 3.8 Å per Cα-Cα distance across a single peptide bond). In beta sheets, the average Cα_1_-Cα_4_ distance is over 10 Å. After examining many structures, we observed that four-residue segments with distance of about 9 Å are L-shaped with three extended residues and the fourth residue hooking left or right. Structures at 8 Å are shaped like turns but wider and shallower than turns with Cα-Cα distances less than 7 Å.

All of our 18 clusters have a median Cα_1_-Cα_4_ distance of 8.0 Å or less, but several of them have significant density above the canonical distance cutoff of 7 Å, especially AB2, dD, dN, and pG. By contrast, the majority of segments in the *Other* category have Cα_1_-Cα_4_ distances of 9 Å or more and are not turns at all but rather extended structures. The median values for the Cα_1_-Cα_4_ distance for each of our clusters are provided in Table 3. We produced

Ramachandran plots for all 18 clusters and the *Other* category without a Cα_1_-Cα_4_ distance cutoff. An example is provided in Fig 8 for AD turns. The rest are provided in S2 Supporting Information. In these plots, we show the Ramachandran distribution of residues 2 and 3 for 4-residue segments with Cα_1_-Cα_4_ distance ≤ 7.0 Å in blue and > 7.0 Å in red, as well as histograms of the distances of each group separately and together. Fig 8 indicates that for AD turns, the distance rises above 7.0 Å in the region where φ_2_≤ −90° and ψ_2_>0°.

**Fig 8.**
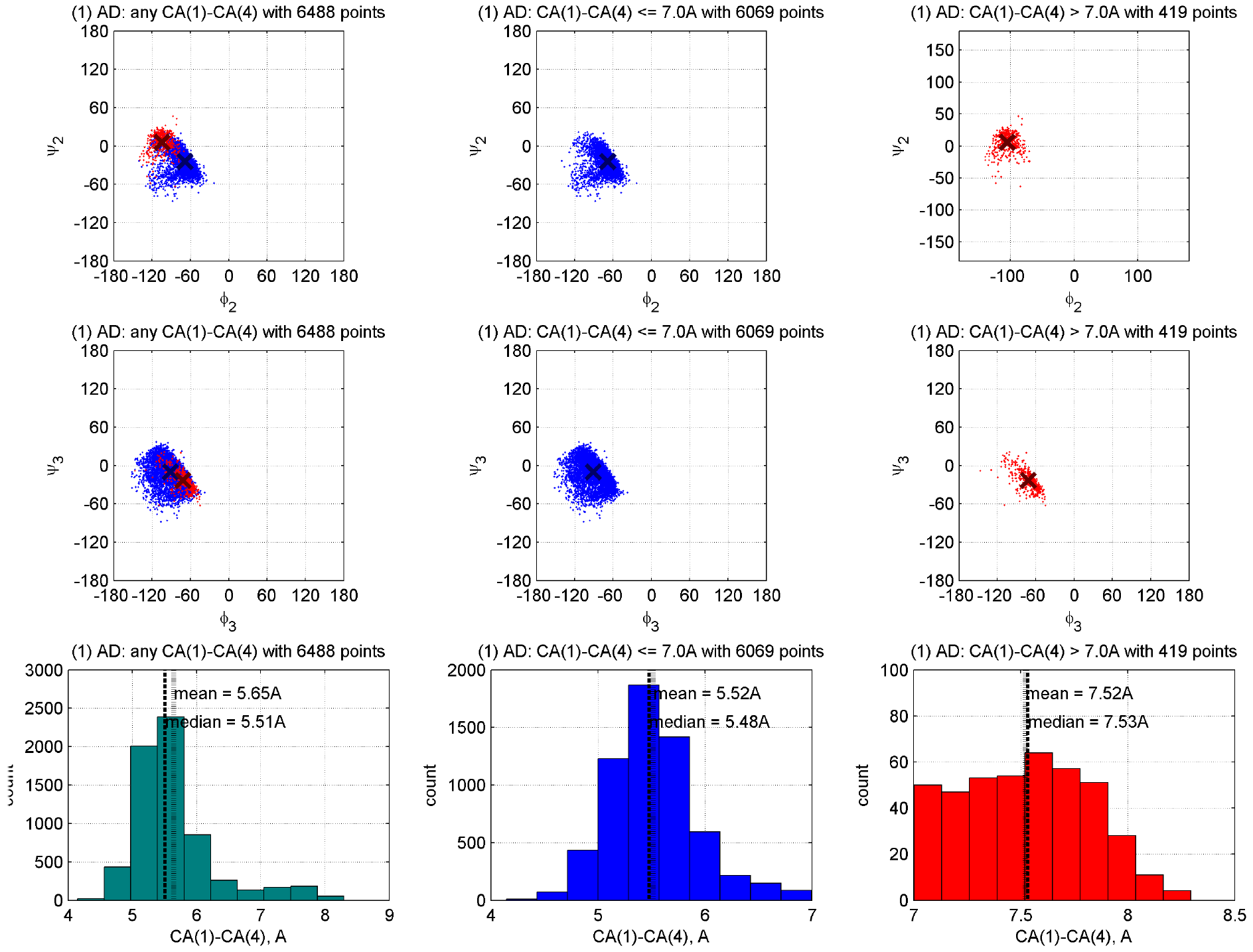
Ramachandran distributions for residues 2 and 3 of AD turns with no Cα_1_-Cα_4_ distance cutoff. In the left column, all points are plotted with those with Cα_1_-Cα_4_ distance ≤ 7.0 Å in blue and Cα_1_-Cα_4_ distance > 7.0 Å in red. Mean (φ, ψ) values are denoted with dark blue and red crosses for each group. The distance histogram is provided in the bottom row. In the middle column, only the points with Cα_1_-Cα_4_ distance ≤ 7.0 Å are plotted, and in the last column only those with Cα_1_-Cα_4_ distance > 7.0 Å are plotted. Figures for the other clusters are provided in S2 Supporting Information.

Given these results, we decided to perform DBSCAN clustering without a Cα_1_-Cα_4_ distance cutoff to check whether that there are additional turn-like clusters hiding in our data set. We found several common Ramachandran combinations for residues 2 and 3, such as BB, aB, and BA, but none of these had average Cα_1_-Cα_4_ distances less than 9 Å, and none of them had density below 7 Å (not shown). At different values of DBSCAN parameters eps and minPts, we could find all of our 18 clusters (sometimes merged as in Fig 1). We conclude that our clustering with the 7 Å cutoff located all significant beta turn types in our data set, and that running our beta turn assignment tool, *BetaTurnTool18*, with different cutoffs up to 8.5 Å may be useful in some circumstances (see below).

### A census of loops, turns, and 3_10_ helices

With a high-resolution data set and a new set of beta turn types, we were interested in the frequency and distribution of beta turns in protein loops of various lengths. This kind of information may be useful in loop structure prediction. For this purpose, we cannot exclude beta turns based on poor electron density or multiple conformations. For residues in our set of 1,074 proteins with alternative conformations, we selected a primary conformation with the highest occupancy. This dataset is named *CompleteSet* containing 224,250 residues with reported coordinates (Table 1).

To identify loops, we needed to consider how to treat 3_10_ helices since three-residue 3_10_ helices occur frequently within longer loops and might be considered part of the loop rather than significant elements of regular secondary structures. To investigate this, we examined kernel density estimates of the Ramachandran maps of residues in 3_10_ helices of different lengths (Fig 9). A large majority of 3_10_ helices are very short – 81% are length 3 and 12% are length 4 (Table 4). 3_10_ helices of length 1 or 2 are impossible in DSSP. The kernel density estimates demonstrate an interesting phenomenon. At positions 1, 2, and 3 of the length-3 3_10_ helices, the mode in the φ-ψ density moves progressively upwards and leftwards: φ,ψ = [−57°,−37°], [−67°,−17°], and [−93°,0°]. This is roughly equivalent to two consecutive AD turns with the middle residue in an intermediate position between A and D. The same is true of length-4 3_10_ helices. For the longer 3_10_ helices, the last residue is always clearly in a delta position [−90°,0°] and the first residue is clearly alpha [−60°,−30°], while the intervening residues are in between, sometimes with points spanning both populations. This is consistent with the early analysis of beta turns by Lewis et al. that defined Type I turns as [φ_2_,ψ_2_ = −60°,−30°; φ_3_,ψ_3_ = −90°,0°] and Type III turns as [φ_2_,ψ_2_ = −60°,−30°; φ_3_,ψ_3_ = −60°,−30°] [11], which were later identified by Richardson as mostly restricted to 3_İ0_ helices [2]. The pattern we observe is slightly more complicated with the modal value shifting from A to D along the 3_10_ helix. The exact modal φ,ψ values for 3_10_ helices of varying length are reported in Table 4. Similar distributions of φ,ψ were found by Pal et al. for short 3_10_ helices [12].

**Fig 9.**
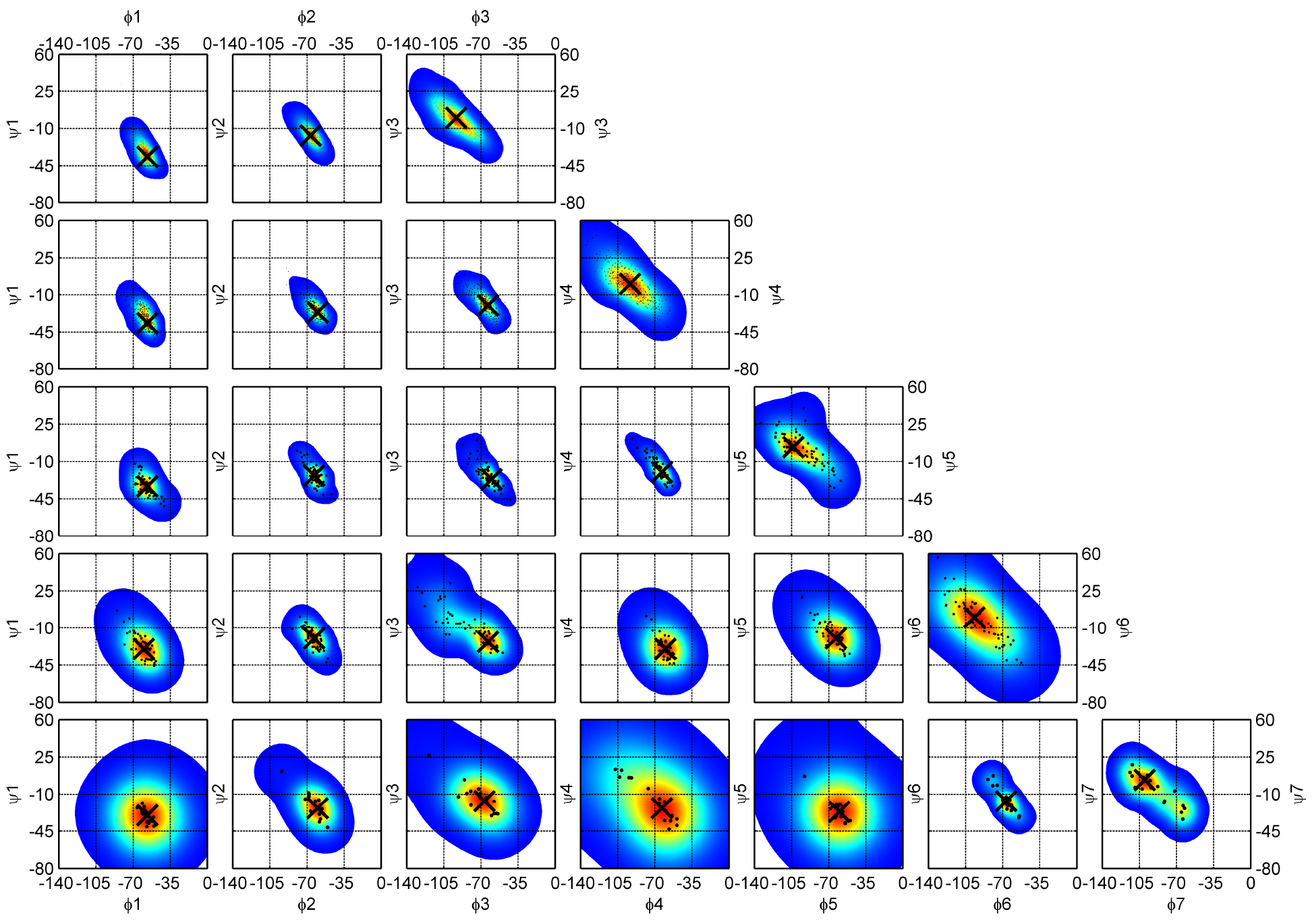
Ramachandran maps for 3_10_ helices of increasing length, *n* from 3 (DSSP code GGG, top row) to 7 (GGGGGGG, bottom row). Optimized kernel density estimates overlaid with *(φ,ψ)* sample scatter plots are drawn for each 3_10_ helix residue starting from the first left position, 1 (first column) to the last right position, 7 (last column). Modes (peaks) for each 2D distribution are shown with a large black cross. The first (1) and last (n) residue conformations of 3_10_ helices are *A* (−64°, −23°) and *D* (−92°, −1°) conformations respectively. The intervening residues are in between, sometimes with conformations spanning both populations. The exact values for (φ,ψ) modes of 3_10_ helices are reported in Table 4.

**Table 4.** Modal values for backbone dihedral angles of 3_10_ helices of lengths from 3 to 7.

| Length | Count | Perc. | $\phi_1, \psi_1$ | $\phi_2, \psi_2$ | $\phi_3, \psi_3$ | $\phi_4, \psi_4$ | $\phi_5, \psi_5$ | $\phi_6, \psi_6$ | $\phi_7, \psi_7$ |
| --- | --- | --- | --- | --- | --- | --- | --- | --- | --- |
| 3 | 2,313 | 80.7 | -57,-37 | -67,-17 | -93, 0 |  |  |  |  |
| 4 | 347 | 12.1 | -57,-37 | -60,-27 | -63,-20 | -93, 0 |  |  |  |
| 5 | 90 | 3.1 | -57,-33 | -63,-23 | -63,-23 | -63,-20 | -103, 3 |  |  |
| 6 | 78 | 2.7 | -60,-30 | -67,-17 | -63,-23 | -60,-30 | -63,-20 | -97, 0 |  |
| 7 | 27 | 0.9 | -57,-30 | -60,-23 | -67,-17 | -63,-23 | -60,-27 | -67,-17 | -100, 3 |
| $\geq 8$ | 12 | 0.4 | - | - | - | - | - | - | - |
| **Total** | 2,867 | 100.0 |  |  |  |  |  |  |  |

We were interested in whether including 3_10_ helices would alter our clusters of beta turns, so we ran our clustering protocol for 3 cases: 1) without GGG (as defined by DSSP) (as already presented above); 2) with GGG; and 3) with both GGG and GGGG. There are no major changes in the clusters except for the AD cluster (Fig C **in S1 Supporting Information**). With the inclusion of GGG and/or GGGG, there are significantly more beta turns in the AD cluster overall. The distribution of φ_3_,ψ_3_ in the AD cluster changes with the inclusion of GGG; a second peak forms near φ_3_,ψ_3_ = [−70°, −20°]. However, we refrained from forming an AA cluster, since there is no significant variation in the amino-acid profiles of the AD and AD/AA clusters in the three cases (Fig D **in S1 Supporting Information**) and the differences in dihedral angles are only about 20° in φ_3_ and ψ_3_.

Given these data, we established two criteria with regard to 3_10_ helices for our definition of protein loops. First, short 3_10_ helices often directly abut alpha helices in DSSP, and represent distortions detected in the *i, i+4* hydrogen bonding pattern of the alpha helix [40]; therefore we exclude such 3_10_ helices immediately adjacent to alpha helices from loops. For example, a continuous fragment with DSSP code (H=helix; G=3_10_ helix; T=turn; C=coil (other)), HHHHGGGG**CCTTCCC**GGGHHHH, has only **CCTTCCC** as a loop while the flanking Gs are still a continuation of an alpha helix region before and after the loop. Second, we treat 3_10_ helices of length 3 not adjacent to alpha helices as loop residues, since they are identified by DSSP with a single internal *i, i+3* hydrogen bond within the loop between the residue that precedes the 3_10_ helix and the last residue of the 3_10_ helix. These short 3_10_ helices are common in long loops and may productively be considered part of the loop rather than an intervening element of regular secondary structure. They are not likely to contribute more than one amino acid to the hydrophobic core, and in many cases not even that. Indeed, in some definitions of secondary structure, a 3_10_ helix requires two or more hydrogen bonds [6] and these three-residue 3_10_ helices would not be identified as such. For example, **GGG** in HHHHCCCC**GGG**CCCCHHHH is still considered a part of the loop with DSSP designation CCCC**GGG**CCCC. We define longer 3_10_ helices (≥4) as elements of secondary structure, which therefore separate their neighboring segments into two loop regions. For instance, GGGGG in HHHHCCCC**GGGGG**CTTCHHHH produces two loops, CCCC and CTTC. To contain a beta turn, a loop must be at least two residues long.

With this loop definition, in *CompleteSet* we detected 21,454 turns with the canonical 7 Å Cα_1_-Cα_4_ cutoff (65% more than in *RefinedSet)* and assigned them to the closest turn type in Table 2 with the distance measure in the seven-dimensional dihedral angle space and the distance cutoff for the *Other* category. Beta turns comprise 29% of residues in the *CompleteSet* and 63% of the residues in loops between regular secondary structures (Table 1). In *CompleteSet*, we located 17,176 loops of length from 1 to 81. The frequency of a loop exponentially decreases with loop length; approximately there is a _10_-fold reduction every time a loop becomes 10 residues longer (Fig 10). 95% of all loops are 1 to 16 residues long. Very long loops are rare; only 0.59% of loops are 30 or more residues long (the 81-residue “loop” is the snow flea antifreeze protein in PDB entry 3BOG (chain A) which has no regular secondary structure as defined by DSSP). The distribution of loop lengths with all 3_10_ helices considered to be regular secondary structure (and not a part of loops) is shown in Fig E **in S1 Supporting Information**.

**Fig 10.**
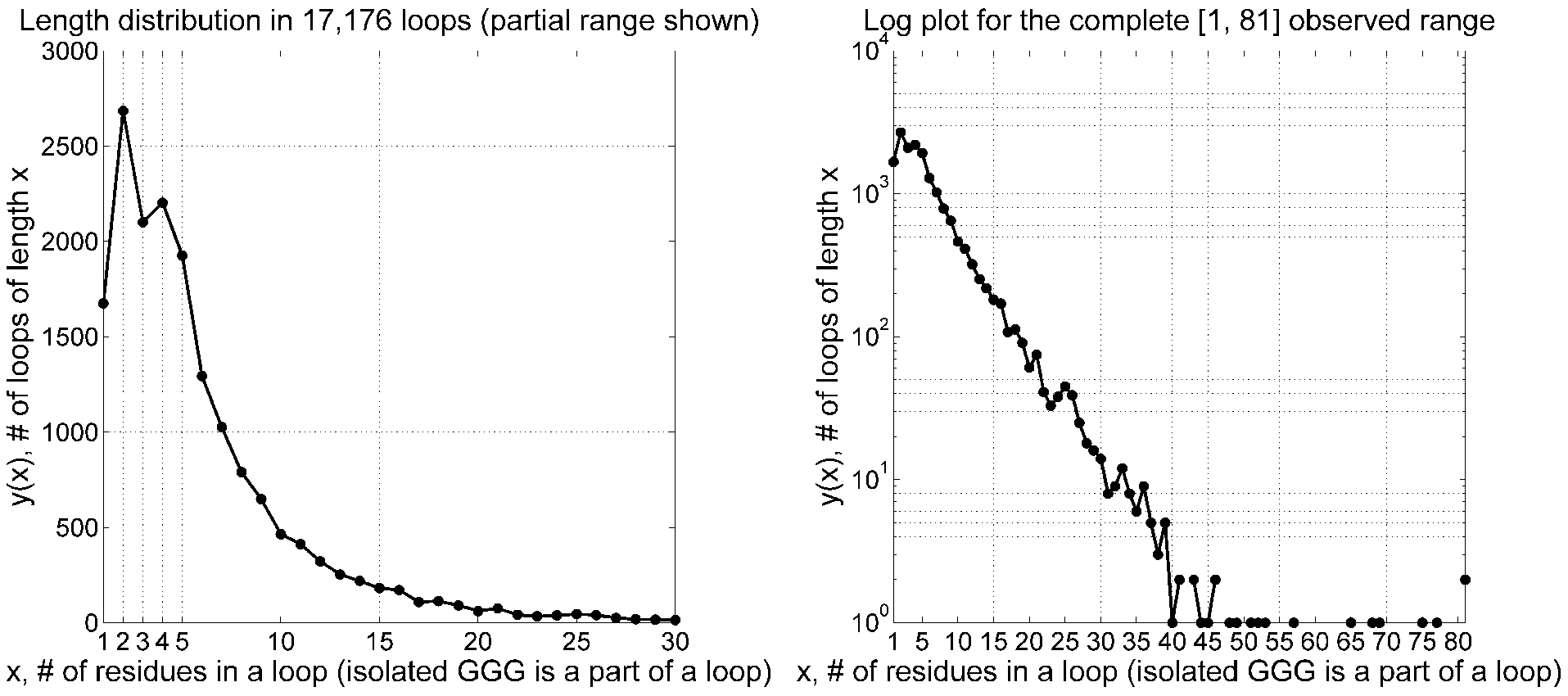
Frequency of loop length in *CompleteSet* of 17,176 loops. Short 3_10_ helices (DSSP of GGG) are considered parts of loops. The frequency of a loop decreases exponentially after the length of 5 with approximately a 10-fold reduction every time a loop adds 10 more residues. Between loop lengths 2 and 5, there is a frequency fluctuation with two peaks at length 2 and 4 observed due primarily to beta turns between two beta-sheet strands. 95% of loops are 1 to 16 residues long. A similar figure for traditional (GGG excluded) loops is provided in Fig E **in S1 Supporting Information**.

We calculated an average number of beta turns identified in each loop of a particular length and estimated ± one standard deviation confidence range (Fig 11). There is an average one beta turn per 4.8 residues. If we consider longer loops with 9 or more residues, there are 11,270 beta turns among 49,299 loop residues, leading to approximately the same value of one beta turn per 4.4 loop residues. The results with 310 helices considered regular secondary structure are shown in Fig F **in S1 Supporting Information**. If 3_10_ helices are considered regular secondary structure and not part of loops, there is an average of one beta turn per 5.5 loop residues for all loops.

**Fig 11.**
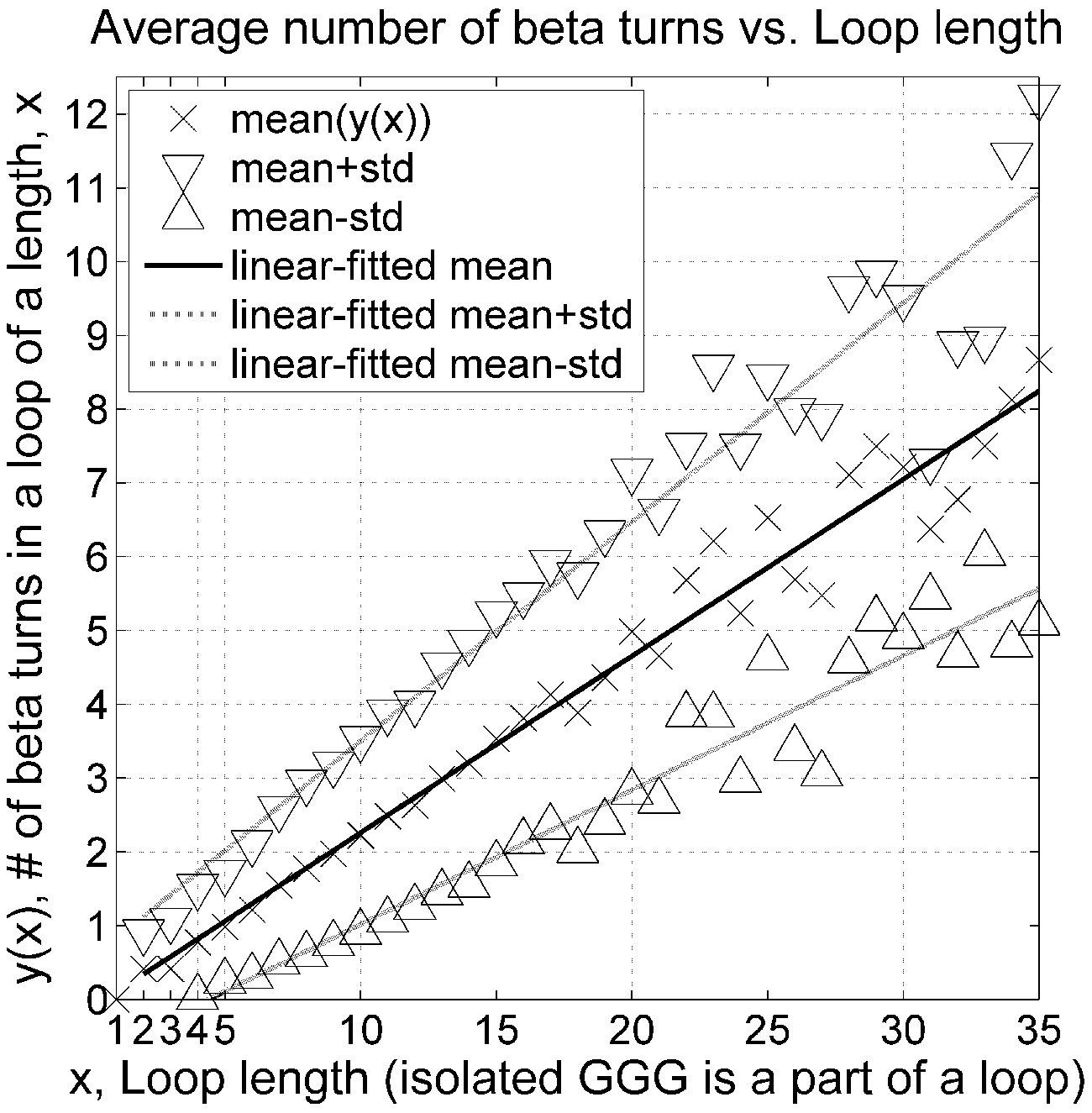
An average number of beta turns in a loop (isolated GGG included in the loops) as a function of loop length. Beta turns are impossible (0 beta turns) in loops with a length of 1 residue because both 2^nd^ and 3^rd^ residues of a beta turn have to be in a loop region. For every 4.8 loop residues, there is on average a single beta turn. A similar figure for traditional loops (GGG excluded) is provided in Fig F in S1 Supporting Information.

A total of 64% (Table 1) of beta turns overlap other beta turns by one or more residues. 310 helices of length 3 plus the preceding residue usually comprise two overlapping beta turns. In addition, other beta turn types can also combine to produce immediately overlapping, consecutive beta turns. We define double turns as a set of 5 residues with residues 1-2-3-4 and

2-3-4-5 forming individual beta turns. These double turns occur most frequently; we found 5,372 double turns having 82 combinations of our 18 turn types (Table 5). The most common double turn (62%) is AD-AD made of two consecutive AD turns, which are the most prevalent turn type in the data set (49%, Table 2). The 10 most frequent turn type combinations account for 90% of all double turns.

**Table 5.** Statistics for 10 most common double turn type combinations in *CompleteSet*. Double turns overlap by three residues (i.e., 1-2-3-4 and 2-3-4-5).

| Rank | Double turn type | Count | Perc. | Cumulative Perc. |
| --- | --- | --- | --- | --- |
| 1 | AD-AD | 3,307 | 61.6 | 61.6 |
| 2 | AD-AZ | 470 | 8.7 | 70.3 |
| 3 | AD-AB2 | 223 | 4.2 | 74.5 |
| 4 | Pa-ad | 199 | 3.7 | 78.2 |
| 5 | AD-AB1 | 137 | 2.6 | 80.7 |
| 6 | AD-AG | 121 | 2.3 | 82.9 |
| 7 | pD-AD | 119 | 2.2 | 85.2 |
| 8 | Pd_dD | 103 | 1.9 | 87.1 |
| 9 | Pd_dN | 84 | 1.6 | 88.7 |
| 10 | PcisD_cisDA | 76 | 1.4 | 90.1 |
|  | other | 533 | 9.9 | 100.0 |
|  | **All** | **5,372** | **100%** | **100%** |

For 1,000 triple turns (Table 6) defined as a stretch of 6 residues with turns at positions 1-2-3-4, 2-3-4-5, and 3-4-5-6, the 7 most frequent combinations out of the observed 65 comprise 82% of triple turns. As for single and double turns, AD-AD-AD is again the most common triple turn, representing half of all cases. Of course turns can overlap by one or two residues (e.g. 1-2-3-4 and 3-4-5-6) but we did not study this separately.

**Table 6.** Seven most common triple turn types in *CompleteSet* among 1,000 observed total. A triple turn is formed by 6 residues with 1-2-3-4, 2-3-4-5, and 3-4-5-6 positions.

| Rank | Triple turn type | Count | Perc. | Cumulative Perc. |
| --- | --- | --- | --- | --- |
| 1 | AD-AD-AD | 486 | 48.6 | 48.6 |
| 2 | AD-AD-AZ | 143 | 14.3 | 62.9 |
| 3 | AD-AD-AB2 | 68 | 6.8 | 69.7 |
| 4 | AD-AD-AG | 37 | 3.7 | 73.4 |
| 5 | Pd_dN_AD | 34 | 3.4 | 76.8 |
| 6 | AD_AD_AB1 | 29 | 2.9 | 79.7 |
| 7 | PcisD_cisDA_AD | 20 | 2.0 | 81.7 |
|  | other | 183 | 18.3 | 100.0 |
|  | **All** | **1,000** | **100%** | **100%** |

### Turn library and software

We have compiled *BetaTurnLib18* files that contain data on all 18 established beta-turn types in S1 Text (tab-delimited text format) and S1 Table (Excel format). For each turn type we provide the following information: the absolute number of dataset points and percentage for each type cluster, the older nomenclature for existing types, our new nomenclature, median and mean CαrCα_4_ distances (both with and without the 7 Å cutoff), medoids and modes for dihedral angles pulled from the *RefinedSet* with the following information: <ω_2_, ϕ_2_, ψ_2_, ω_3_, ϕ_3_, ψ_3_, ω_4_>, <aa_1_, aa_2_, aa_3_, aa_4_>, <sec_1_, sec_2_, sec_3_, sec_4_>, res_id(1), chain_id and pdb_id. Our library, *BetaTurnLib18* is freely available to be incorporated into any third-party software for detection, type prediction, and modeling of beta turns in proteins.

In addition, we provide Python software, *BetaTurnTool18* supported on Linux, Mac, and Windows platforms coded with back-compatible Python 2. It reads a PDB-formatted or mmCIF-formatted coordinate file of a protein structure, automatically runs DSSP to assign secondary structure in the input file, and parses our turn library file. With this information available, the Python script detects beta-turn positions and types. Beta-turn location is determined by verifying chain connectivity of each four-residue stretch, satisfying the default 7.0 Å Cα_1_-Cα_4_ distance constraint calculated from the Cartesian coordinates and checking the secondary structure of the 2^nd^ an 3^rd^ residues assigned by the DSSP program [10]. The program also allows the user to set a different Cα_1_-Cα_4_ distance cutoff, which may be useful for some beta turn types that possess a broader distribution of Cα_1_-Cα_4_ distances (Fig 7). In addition, *BetaTurnTool18* has two options to search for beta turns within loops either with or without inclusion of isolated 310 helices.

Since some beta turns are similar to more than one of our beta turn types, the first and second most probable turn types are reported based on the distance to the first and second closest medoids calculated with the same distance metric as used in the clustering, unless the closest medoid is greater than 0.2359 units in our distance metric, in which case the turn is reported as *Other*. We estimate confidence levels for the first and second most probable types, which sum up to 1, as follows:

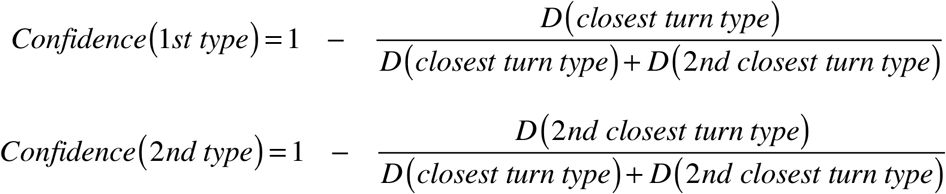

Our script prints these confidence levels as integers in the [0, 9] range. 0 and 9 correspond to the [0, 10)% and [90, 100]% confidence intervals respectively. A border-case flag (“+”) is returned when the integer confidence level is 6 or below, and a “.” (dot) is returned when it 7 or higher. In border cases we recommend considering both turn types for modeling. Our tool accounts for chain breaks in coordinates when it prints a single-character alignment of the complete amino acid sequence, its DSSP secondary structure, turn types, integer confidence levels, and border-case flags. Downloads and complete documentation of our tool or library are available at dunbrack.fccc.edu/betaturn and github.com/sh-maxim/BetaTurn18.

## Discussion

From a set of 13,030 beta turns identified in 1,074 proteins of resolution 1.2 Å or better, we have identified 18 beta turn types, compared to the 8 beta turn types in common use (I, I’, II, II’, VIa1, VIa2, VIb, and VIII). Our nomenclature is based on positions in the Ramachandran map of residues 2 and 3. Six of the turn types are essentially new: dD, dN, Dd, pG, cisDA, and cisDP. Two new types of turns with left-handed residue 2 and right-handed residue 3 (“dD” and “dN”), and the new complementary turn type with right-handed residue 2 and left-handed residue 3 (“Dd”). There is a new turn type “pG” and two new turn types with a cis peptide bond at position 2: “cisDA” and “cisDP.”

Four of the existing turn type remain essentially unchanged but have been renamed: Type I is now “AD” and its complement Type I’ is now “ad”; Type II’ is now “pD”; and Type VIa1 with a cis peptide bond at residue 3 is now “PcisD”. Eight turn types are subtypes of the classical turn types: Type VIII turns have been replaced with 4 subtypes (“AB1”, “AB2”, “AZ”, and “AG”) with distinct peaks in the density in the Ramachandran map of residue 3 and distinct residue preferences at one or more of the 4 turn positions. Type II turns now consist of two subtypes, “Pd” and “Pa”, defined by the presence or absence of Gly at residue 3 respectively. Type VIb turn types have been split into two: BcisP (close to the original definition) and PcisP with a change in ϕ_2_. Turns of Type VIa2 with a cis peptide bond at residue 3, ϕ_2_, ψ_2_=[−120°,120°] and ϕ_3_,ψ_3_=[−_60_°,_0_°], were too spread-out and contained fewer than 25 points in *RefinedSet* (0.2%), too few to form a cluster; they were left in the *Other* group.

Our decisions in clustering were based on two criteria: keeping the total number of turn types at a minimal practical level for existing and future applications; and justifying new cluster formation either by a significant distance from other formed clusters or uniqueness of a four-residue turn profile. Thus, we abandoned formation of a new cluster if its amino acid turn profile was not unique enough or its conformation spread could be justified by continuous, low-energy torsion angle variation. After running DBSCAN and subdividing some clusters, 2.4% of the turns were left as *Other*. We conclude that the 18 types cover 98% of turn conformations and that we have not overlooked any major turn conformations. We have also taken a census of beta turns in loops in our data set and determined that there is one turn per 4.8 residues, regardless of loop length. Many turns overlap other turns, such that 63% of residues in loops are in beta turns.

We provide a tab-delimited library file, *BetaTurnLib18* of the new 18 turn types. For each type, we list proposed and existing names, percentage, and geometrical conformation expressed with torsion angles of a real dataset sample. For each cluster, two verified sample conformations with good electron density are given: closest to a mode and closest to a medoid. This library may be used in third-party software for modeling and prediction of beta turns. In addition, we prepared an easy-to-use Python tool, *BetaTurnTool18* (supported on Linux, Mac and Windows platforms) that reads a PDB-formatted or mmCIF formatted coordinate file of a protein structure as input. The tool assigns secondary structure with included redistributable DSSP program, uploads our turn library, and finally locates turns and identifies their types. The output includes the first and second most probable types with assigned confidence levels for each detected beta turn.

Beta turns in proteins have been studied for 50 years and many classification schemes have been presented. Ultimately a consensus has arisen consisting of 8 turn types: I, I’, II, II’, VIa1, VIa2, VIb, VIII, and a catch-all *“Other”* turn type IV. We have found that some of these turn types represent more than one mode in the density and should be subdivided into further turn types. One of the more interesting results of the new clustering are the turn types that represent combinations of right-handed helical and left-handed helical conformations of residues 2 and 3. We find both combinations: left-right (types dN and dD) and right-left (Dd). These are *not* the same as the type V and V’ turns defined by Lewis et al. [11], which had dihedrals of (−80°,+80°) and (+80°,−80°) for type V and (+80°,−80°) and (−80°,+80°) for type V’. The dihedral angle modes for turn types dN, dD, and Dd are substantially different from the Type V and V’ values (Table 2).

Pal and Chakrabarti performed an extensive analysis of cis peptide bonds in proteins, and in particular they classified turns with cis peptide bonds at position 3 [41]. They divided the turns based on the presence of hydrogen bonds rather than the dihedral angle values of connecting the Cα_1_ and Cα__4__ atoms as we have done. This resulted in a set of seven turn types, with different dihedral angle values provided depending on whether residues 2 and 3 were both proline (Pro-Pro), both not proline (Xnp-Xnp) or the more common Xnp-Pro type. Because their data set was very small (147 proteins), only 4 of the types had more than 10 examples: VIa1, VIa2, VIb1, and VIb2. The others are VIb3, VIc, and VId.

Their VIa1 and VIa2 types are distinguished by the presence or absence respectively of a hydrogen bond between the carbonyl oxygen of residue 1 and the NH of residue 4. The VIa2 turns have a more negative value of ϕ_2_ than VIa1 turns for both Xnp-Pro and Xnp-Xnp cases. We have 125 PcisD turns which are equivalent to VIa1 turns, two of which are Xnp-Xnp and 15 of which are Pro-Pro turns. Our Xnp-Xnp and Pro-Pro PcisD turns are closer to the medoid of PcisD than they are to the values quoted by Pal and Chakrabarti. As described above, even in our much larger data set, we only have 15 beta turns within ± 30° of the VIa2 type (as defined by Hutchinson and Thornton [17]), all of which are of the Xnp-Pro type. They are distributed within a 50° by 50° region in ϕ_2_, ψ_2_. These are classified as *Other* in our set of turn types.

The VIb1 and VIb2 beta turns defined by Pal and Chakrabarti are distinguished by the absence or presence of a hydrogen bond of the residues immediately before and after the beta turn. The VIb3 turn has a CH-O hydrogen bond between residues 2 and 3 and a ϕ_2_ near 180°. We find the Xnp-Pro VIb1 and VIb2 turns of Pal and Chakrabarti are well represented in our data. Turns near their VIb1 values are a mix of our BcisP and PcisP clusters; turns near their VIb2 values are all BcisP. We have six Pro-Pro turns in PcisP, three closer to their VIb1 and three closer to their VIb2 but all closer to our PcisP medoid. We find only five Xnp-Pro turns close to the dihedral values of VIb3 turns, but they are also closer to our medoid than VIb3. We find no VIc or VId turns in our data set.

Our new clustering will be useful in determining and refining structures with X-ray crystallography, NMR, and cryo-EM methods. Structure determination, especially at low-resolution, is dependent on templates of common conformations and statistical distributions of backbone and side-chain distributions [42]. Loops are particularly difficult to model at low resolution [43] and our clear definitions of beta turns may be useful in refining these elements of protein structures. Long loops are also difficult to model during the process of structure prediction by comparative modeling [44, 45], and better sampling and modeling of beta turns may prove useful. Finally, the interpretation of missense mutations, either inherited or those that arise somatically in cancer, remains challenging [46]. Structural approaches have contributed to solving this problem, and mutations in beta turns may be particularly disruptive to loop conformations [47]. Given the ubiquitous nature of beta turns, the clustering, nomenclature, and identification script we have developed should provide useful tools for protein structure determination, refinement, and prediction programs as well as useful tools for protein structure, function, and evolutionary analysis.

## Methods

### Data set

A dataset of high-resolution 1,074 protein chains was compiled with PISCES [28, 29]. It consists of PDB chains of length 40 or more amino acids, from X-ray structures with resolution 1.2 Å or better, R-factor of 0.2 or better, and 50% or lower inter-chain sequence identity, which is now determined by local HMM-HMM alignment with HHSearch [48]. Their 8-label DSSP secondary structure was established from a large file holding DSSP secondary structure for the entire PDB which was downloaded from the RCSB website. This raw dataset had 232,197 residues in amino acid sequences, and 224,250 residues with known backbone coordinates. From this raw dataset we compiled two datasets: *CompleteSet* and *RefinedSet*. For *CompleteSet* we extracted residues with at least backbone coordinates for N, Cα, and C atoms and chose a primary conformation for each residue. For a single-conformational residue, there is only one conformation available and it was pulled into *CompleteSet*. For a multi-conformational residues, a primary conformation was pulled into *CompleteSet*. The primary residue conformation has the highest occupancies for its atoms (in PDB files, it is predominantly the *‘A’* alternative conformation but not always). *CompleteSet* was used for beta-turn statistical analysis which is representative of the general protein population.

Next we generated *RefinedSet* which is a subset of *CompleteSet. RefinedSet* contains a set of higher-quality residues to generate a set of higher-quality beta turns for clustering to identify reliable beta-turn types. The following steps were taken to generate *RefinedSet*: (1) We skipped all residues with any missing backbone, Cβ, and/or γ heavy atoms; (2) we skipped residues with more than one conformation and required the atom occupancy to be 1.0 for the backbone atoms; (3) we combined the DSSP secondary structure information with a set of PDB coordinates and skipped any residues with undefined DSSP secondary structure. These actions produced *RefinedSet* of a total of 195,332 residues.

We detected beta turns in *RefinedSet* according to the following beta-turn definition. For the clustering procedure, a contiguous four-residue segment is a beta turn if the Cα atoms of the 1^st^ and 4^th^ residues have a distance of ≤ 7.0 Å and the 8-label secondary structure of the 2^nd^ and 3^rd^ residues is different from beta-sheet (E), alpha-helix (H), π-helix (I) and 3_10_-helix (G) according to DSSP [10].

For the census of beta turns within loops in *CompleteSet*, (1) we counted three-residue 3_10_-helices within loops (CCCC**GGG**CCCC) as part of the loop and the residues were allowed to be labeled beta turns; (2) we excluded 3_10_-helices immediately adjacent to alpha helices (HHHHGGG, GGGHHHH, etc.).

### Clustering

For beta-turn conformation clustering, we have to define a distance metric – a discriminative measure of how different two beta-turn conformations are. Our beta turn definition is defined by the Cα(1)-Cα(4) distance. Starting from Cα(1), the Cα(4) atom can be built from standard bond lengths and bond angles and seven dihedral angles: ω(2), ϕ(2), ψ(2), ω(3), ϕ(3), ψ(3), ω(4). Inclusion of two flanking torsion angles: ψ(1) and ϕ(4) created too many sub-clusters with no distinct amino acid signatures (not shown).

Since torsion angles are periodic, we used a metric from directional statistics [49], which is equal to the squared chord length between a pair of torsion angle values placed on the unit circle:

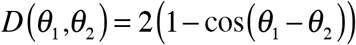

The factor of 2 is unnecessary but we have included it as we did in earlier work on the clustering of conformations of complementarity-determining regions of antibodies [25]. For two beta turns, the distance between them is:

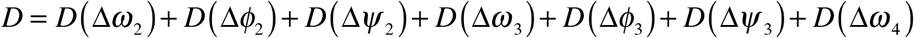

where the differences in angles are taken between the dihedral angles in the two turns.

Clustering of the *RefinedSet* was performed with the DBSCAN algorithm. We manually optimized the *eps* and *minPts* in order to minimize the number of *Other* points and to produce clusters with [ϕ(2), ψ(2)] and [ϕ(3), ψ(3)] restricted to well-known regions of the Ramachandran map for each cluster. This was possible for all the clusters returned by DBSCAN with parameters *eps* = 14° and *minPts* = 25, with the exception of Type I and Type VIII turns. These turns could not readily be separated by DBSCAN since the density for residue 3 is more or less continuous from the alpha to the beta regions of the Ramachandran map. Kernel density estimates of this cluster showed 5 peaks in the density. Three other clusters also showed more than one peak in the density of residue 3. The *k*-medoids algorithm was applied sequentially to these four clusters in order to find clusters associated with each peak in the [ϕ(3), ψ(3)] density.

For each of resulting 18 clusters, we identified and report both medoid and mode of a cluster density in the seven-dimensional ω_2_-ϕ_2_-ψ_2_-ω_3_-ϕ_3_-ψ_3_-ω_4_ dihedral space. The medoid and mode serve as representative conformations for each turn type. We scan a distance matrix of each cluster to find a sample with the most number of neighbors within the smallest radius covering at least 20% of cluster members and at least 20 data points. We approximate a mode with such a turn sample closest to it. The medoid conformation is more sensitive for turn type detection while the mode conformation is more useful in modeling of an unknown protein structure.

### Kernel density estimates

We calculated 1D and 2D kernel density estimates (KDE) for a single circular variable (a single torsion angle) and two circular variables (two torsion angles) respectively with a von Mises kernel which is suitable for circular statistics. For example, for the Ramachandran map distribution, we use a two-dimensional kernel density estimate that we developed for the backbone-dependent rotamer library [50]:

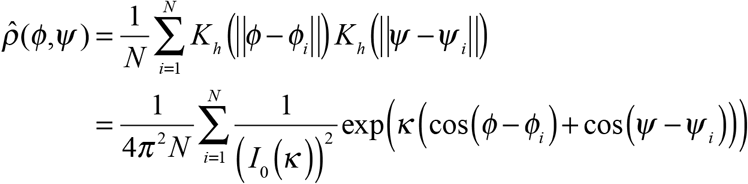

We utilized a 10-fold maximum-likelihood cross-validation to establish the concentration parameter κ for each kernel density estimate.

### Selecting cluster representatives for the beta turn assignment tool

For 36 medoid / mode representatives, we report beta turns satisfying these these conditions: 1) close to medoid / mode by our distance metric, 2) atomic EDIA backbone and side-chain atom electron density better than the median electron density in each cluster, 3) passing a visual inspection of protein geometry and 2Fo-Fc and Fo-Fc electron density maps for any inconsistencies in the placement of backbone and side-chain atoms of the i-1 to *i+4* residues. We manually selected turns that satisfy these criteria.

### Software and Libraries used

Our Python tool for beta-turn type assignment depends on widespread python2 language, numpy [51] and biopython [52] libraries and DSSP software [10, 53]. These are free redistributable software and are easy to download and install. In addition to biopython and numpy libraries we used sklearn [51] and scipy [54] libraries for dataset preparation, residue pruning, and clustering. These libraries allowed PDB parsing, bond and torsion angle calculation, clustering, and scatter plotting. We downloaded electron density maps from European Bioinformatics Institute at www.ebi.ac.uk and ran EDIA software to analyze agreement between turn conformations and their electron density [35].

## Supporting information

S1 Text. BetaTurnLib18 beta-turn library in tab-delimited text format: modes and medoids and other data for all 18 clusters. Version 1.1.

S1 Table. BetaTurnLib18 beta-turn library in Excel format: modes and medoids and other data for all 18 clusters. Version 1.1.

S1 Data Set. A set of 1,074 protein chains used for the beta-turn library compilation. Version 1.1.

---

**S1 Supporting Information**. This file contains supplementary Figures A–F.

**S2 Supporting Information**. This file contains Ramachandran plots for all 18 clusters and *Other* category turns when no Cα_1_-Cα_4_ distance cutoff is used.

**S1 Text.** *BetaTurnLib18* beta-turn library in tab-delimited text format: modes and medoids and other data for all 18 clusters.

**S1 Table.** *BetaTurnLib18* beta-turn library in Excel format: modes and medoids and other data for all 18 clusters.

**Fig A.**
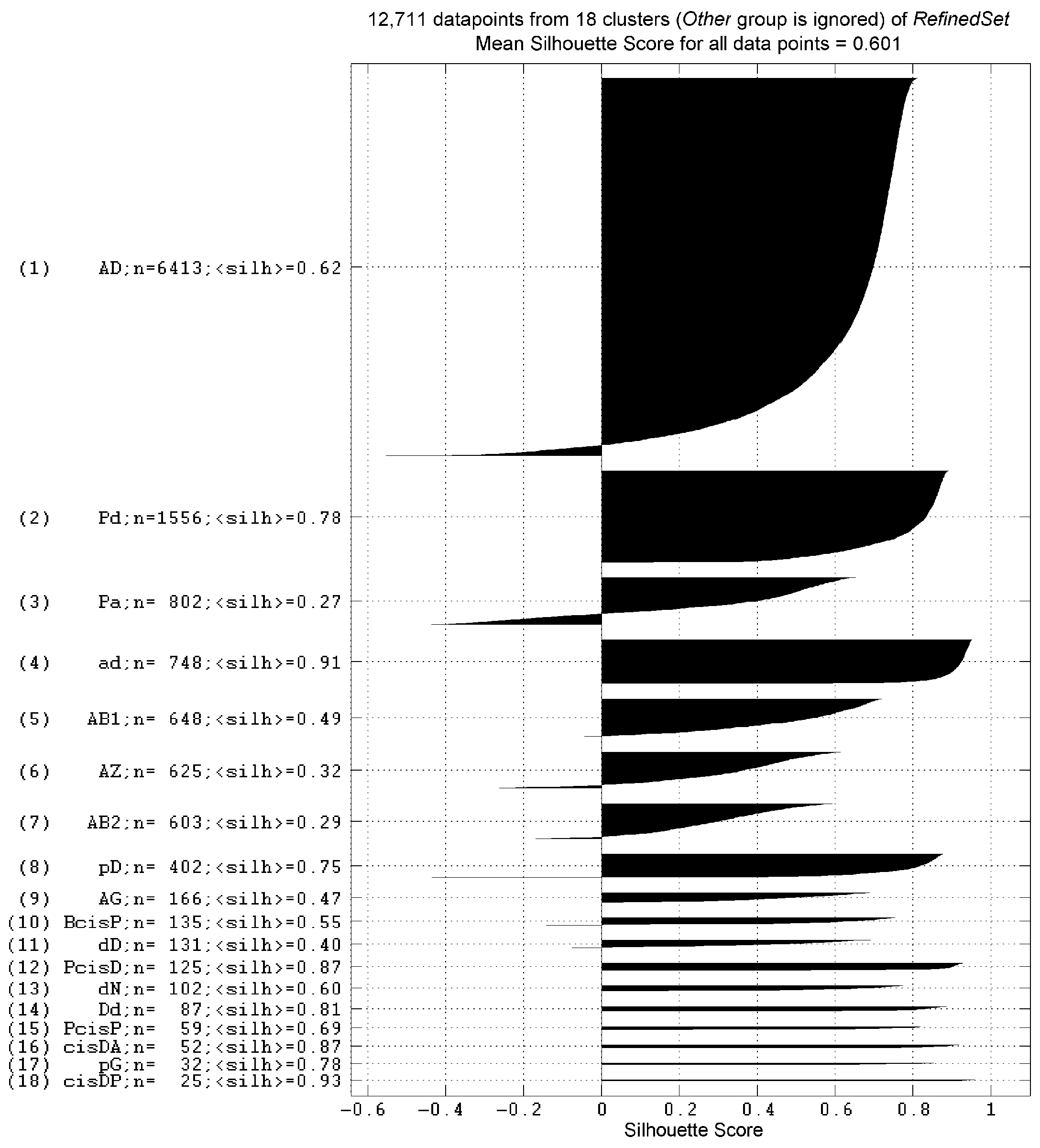
Silhouette scores for all 18 clusters. The overall mean score is 0.601. Some of the clusters produced with k-medoids have lower scores in the (0.3, 0.7) range. DBSCAN failed to produce them in the first place because of their varying density, overlapping and spread-out nature. Some of the clusters have few individual data points with negative (-1.0, 0.0) scores, which occur in some of the subclusters generated by k-medoids. The silhouette score uses a different metric (average distance of a point to the members of a cluster) than k-medoids does (distance of a point to the medoid of a cluster), so that some of the k-medoids clusters have some negative silhouette scores. The silhouette score analysis provides an additional validation of the 18 produced clusters, also supported by distinct peaks in the density generated with a maximum-likelihood optimization and 10-fold cross validation.

**Fig B.**
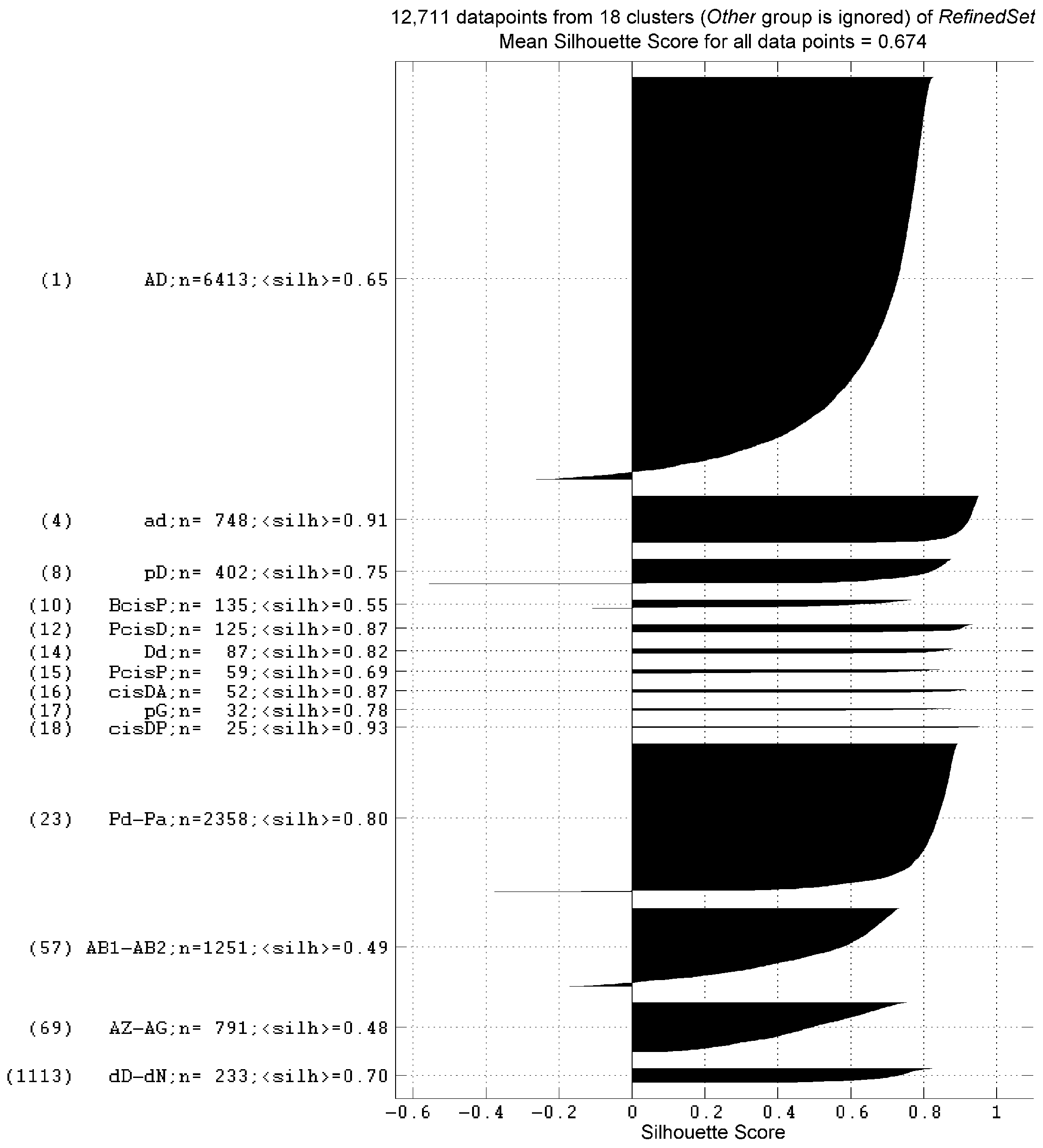
Silhouette scores for 14 clusters after merging subclusters AB1 with AB2, AZ with AG, Pd with Pa, and dD with dN. The mean score of the points after merging is 0.674. The merged subclusters have better silhouette scores than the unmerged subclusters (Suppl. Fig A) but still represent peaks in the density in dihedral angle space and distinct sequence profiles.

**Fig C.**
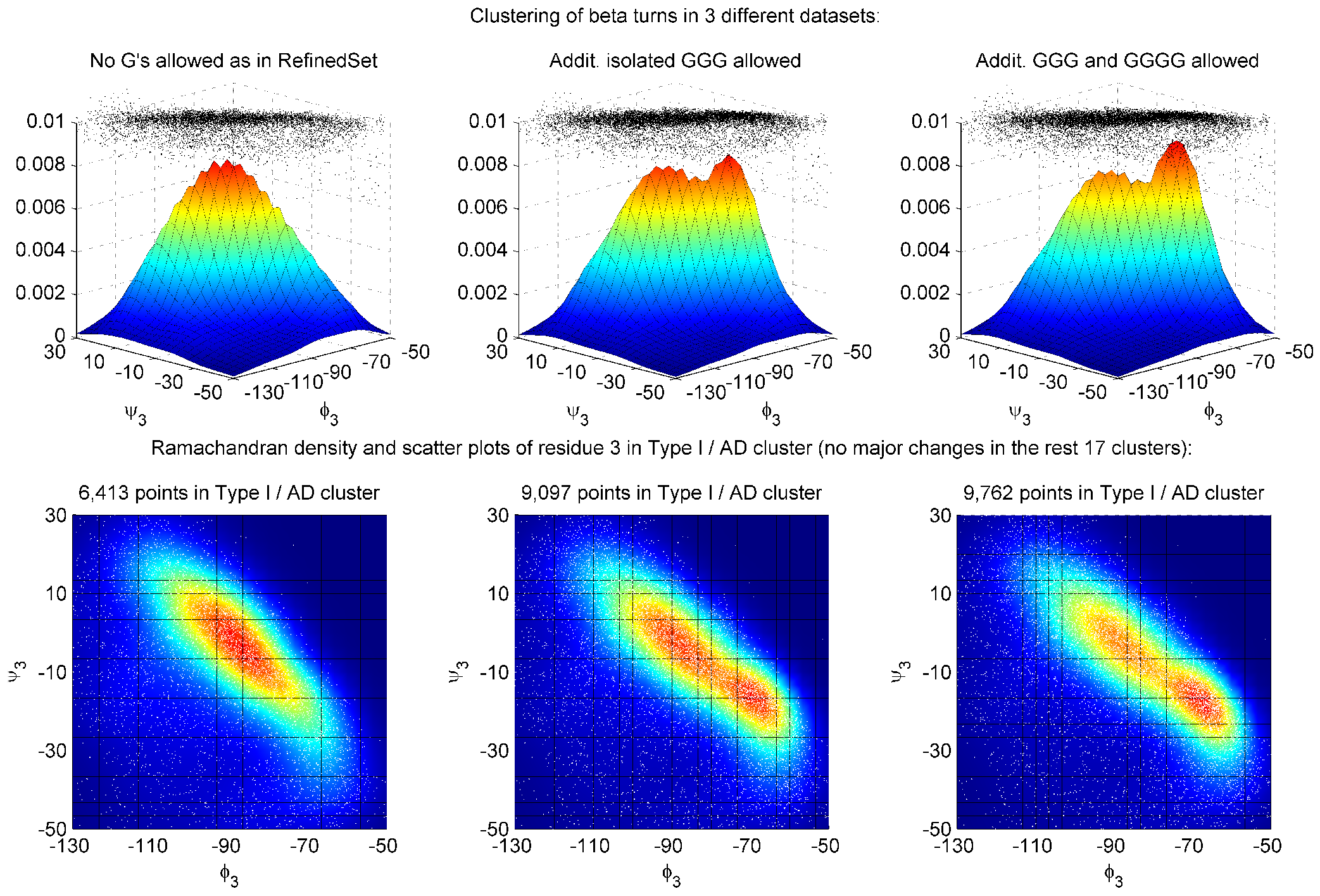
Clustering results for the Type I / AD cluster for 3 cases from left to right: the original *RefinedSet* with no 3io helix (DSSP code G) allowed; the *RefinedSet* with isolated 3-residue 3_10_ helices (GGG) added; and *RefinedSet* with both isolated 3-and 4-residue 3_10_ helices (GGG and GGGG) added. Isolated 3_10_ helices are those not immediately abutting alpha helices. The remaining 17 clusters do not have major changes. The number of beta turns increased by 42% and 52% for the AD cluster overall for GGG and GGG/GGGG respectively relative to no G’s. The second peak near ϕ_3_,ψ_3_ = (−70°,−20°) becomes more prominent: 36% and 38% AA sub-population for GGG and GGG/GGGG vs. 28% for no G’s. The formation of the additional cluster AA was rejected because it did not have significantly different aminoacid profile (Fig D in S1 Supporting Information) and the ϕ_3_,ψ_3_ conformation varies along the natural main diagonal.

**Fig D.**
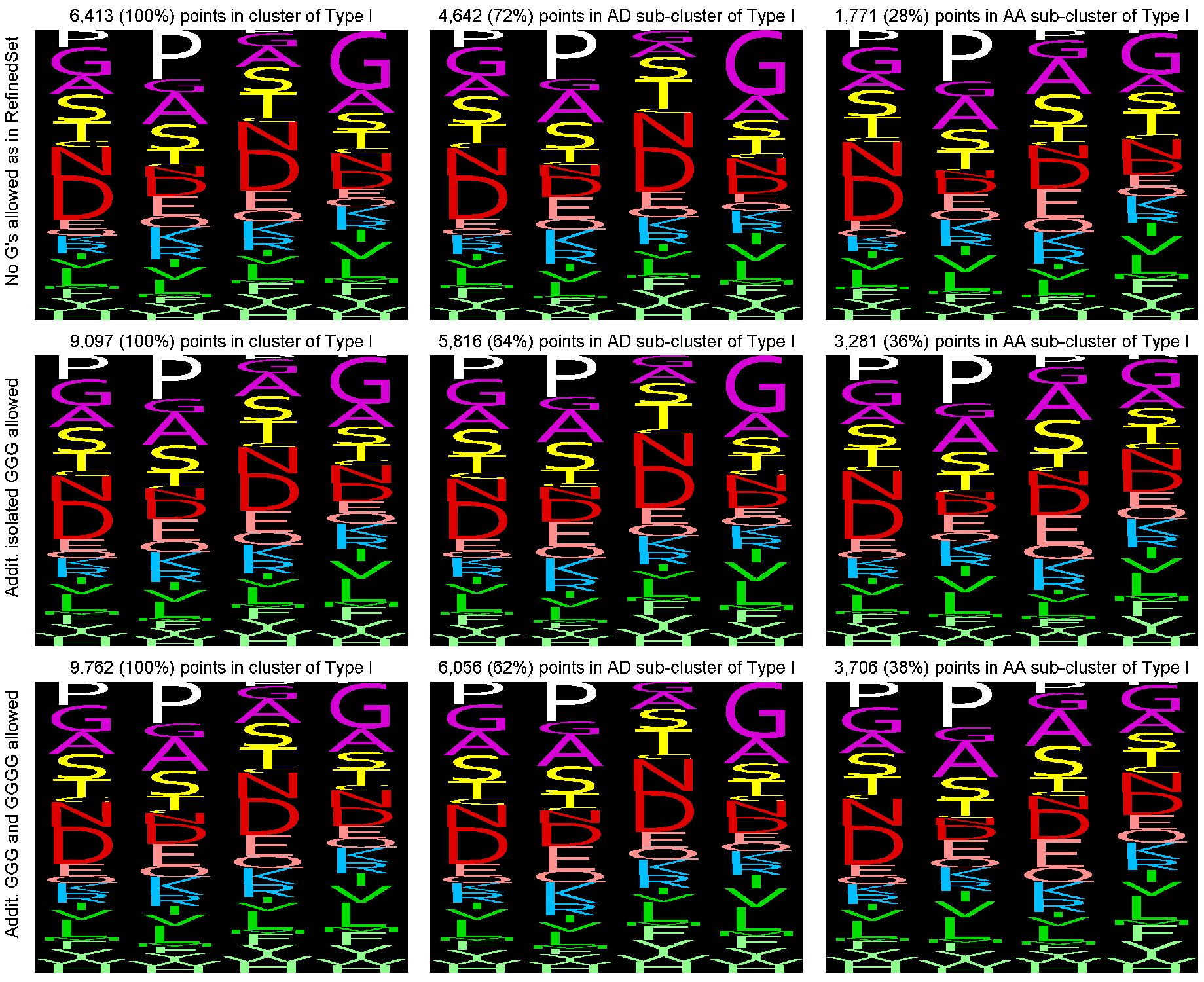
Sequence profiles for clustering with and without 3_10_ helices. First row: *RefinedSet* with no G’s; second row *RefinedSet* with isolated GGG segments added; third row, *RefinedSet* with GGG and GGGG added. First column, the AD+AA cluster; second column, the AD cluster; third column, the AA cluster. The AD / AA sub-clusters populations change from 72% / 28% (no G’s) to 64% / 36% (+GGG) and 62% / 38% (+GGG and GGGG). The formation of the additional cluster AA was rejected because it did not have significantly different aminoacid profile.

**Fig E.**
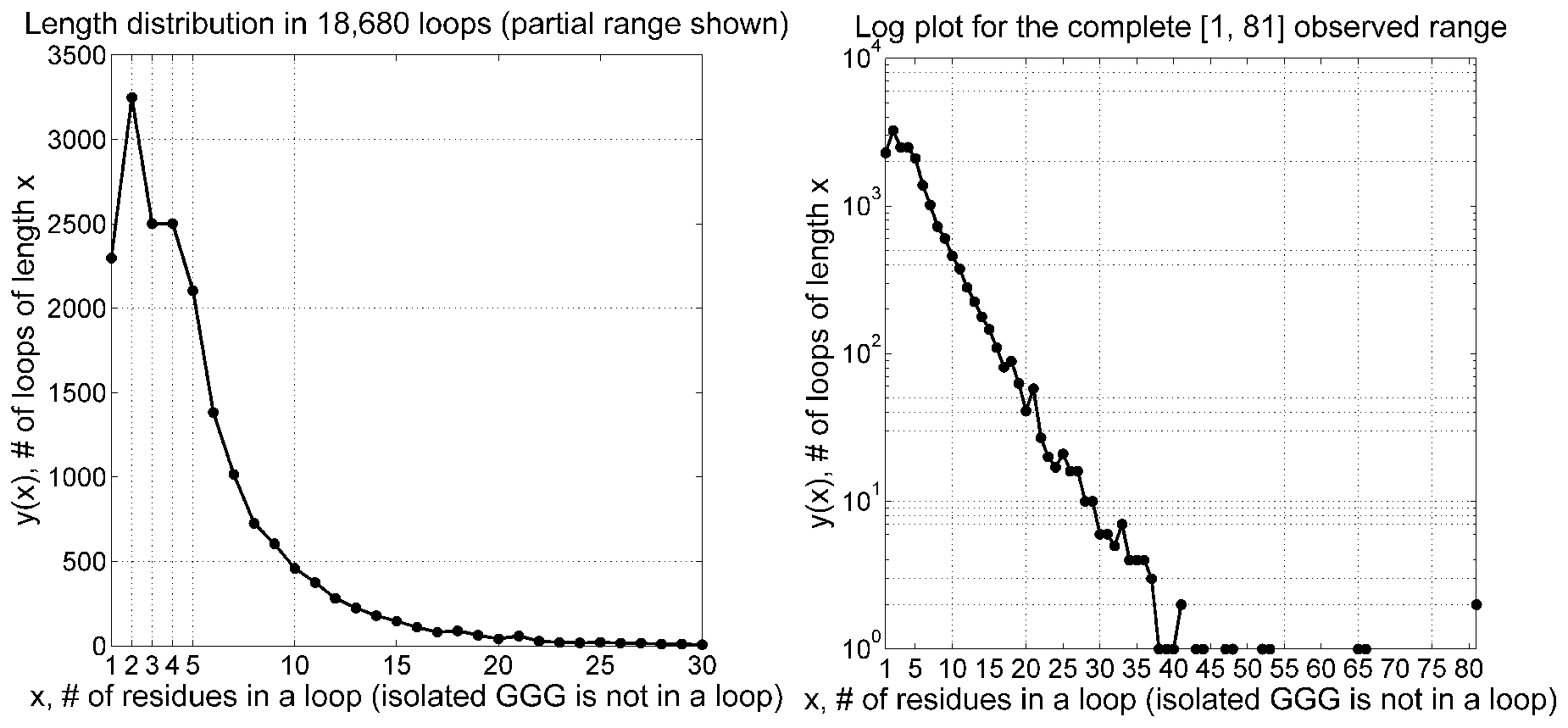
Frequency of loop length in *CompleteSet* of 18,680 loops with all 3_10_ helices excluded from loops. The exclusion of GGG in the loop definition increases the total number of loops by 9% from 17,176 to 18,680, because some loops are split by 3-residue 3_10_ helices. The frequency of a loop still decreases exponentially after the length of 5 with approximately a 10-fold reduction every time a loop adds 10 more residues. 95% of loops are 1 to 13 residues long. Very long loops are rare, only 0.29% of loops are 30 or more residues long.

**Fig F.**
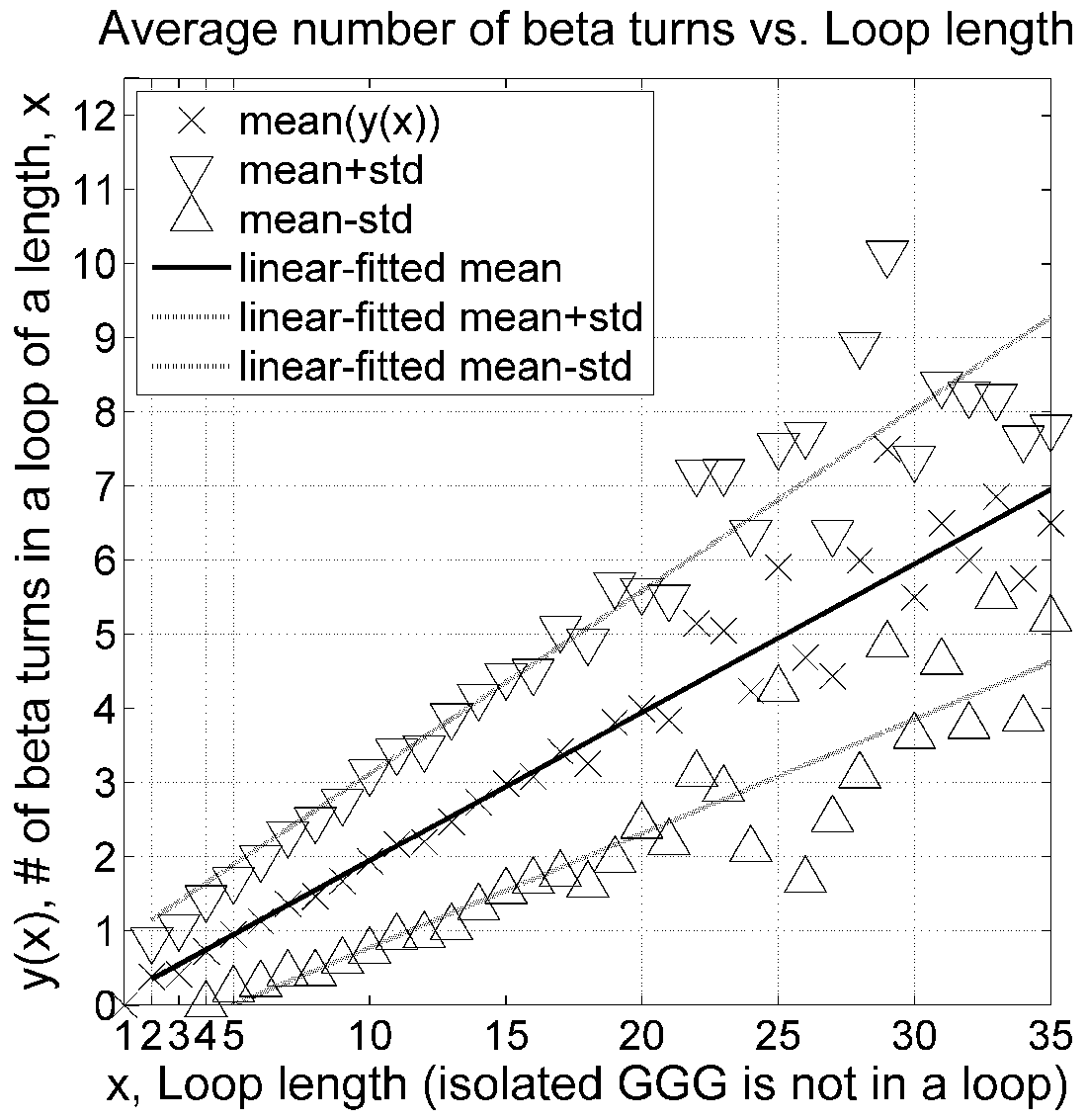
Average number of beta turns in loops as a function of loop length when all 3_10_ helices are not considered part of loops. For every 5.5 loop residues, there is on average a single beta turn.

**S2 Supporting Information.** This file contains Ramachandran plots for all 18 clusters and *Other* turns when no Cα_1_-Cα_4_ distance cutoff is used. Points in blue have Cα_1_-Cα_4_ distance ≤ 7.0 Å; points in red have Cα_1_-Cα_4_ > 7 Å. The first column shows all points that can be assigned to the cluster by the dihedral angle metric regardless of Cα_1_ −Cα_4_ distance. The second column shows data for those points with Cα_1_ −Cα_4_ ≤ 7.0 Å. The third column shows data for those points with Cα_1_-Cα_4_ > 7 Å. The histograms in the bottom row show the distribution of Cα_1_-Cα_4_ distances for each group. There is one page for each cluster and for the *Other* group. Four-residue segments with Cα_1_-Cα_4_ up to about 8 Å are shaped like turns. Those above 7 Å are wider and shallower than classical beta turns (Cα_1_-Cα_4_ ≤ 7.0 Å). Segments with Cα_1_-Cα_4_ distance around 9 Å are L-shaped. Those at about 10 Å are significantly extended and similar to segments of beta sheet strands. Most 4-residue segments in the *Other* category are not shaped like turns since their Cα_1_-Cα_4_ distance is above 9 Å.

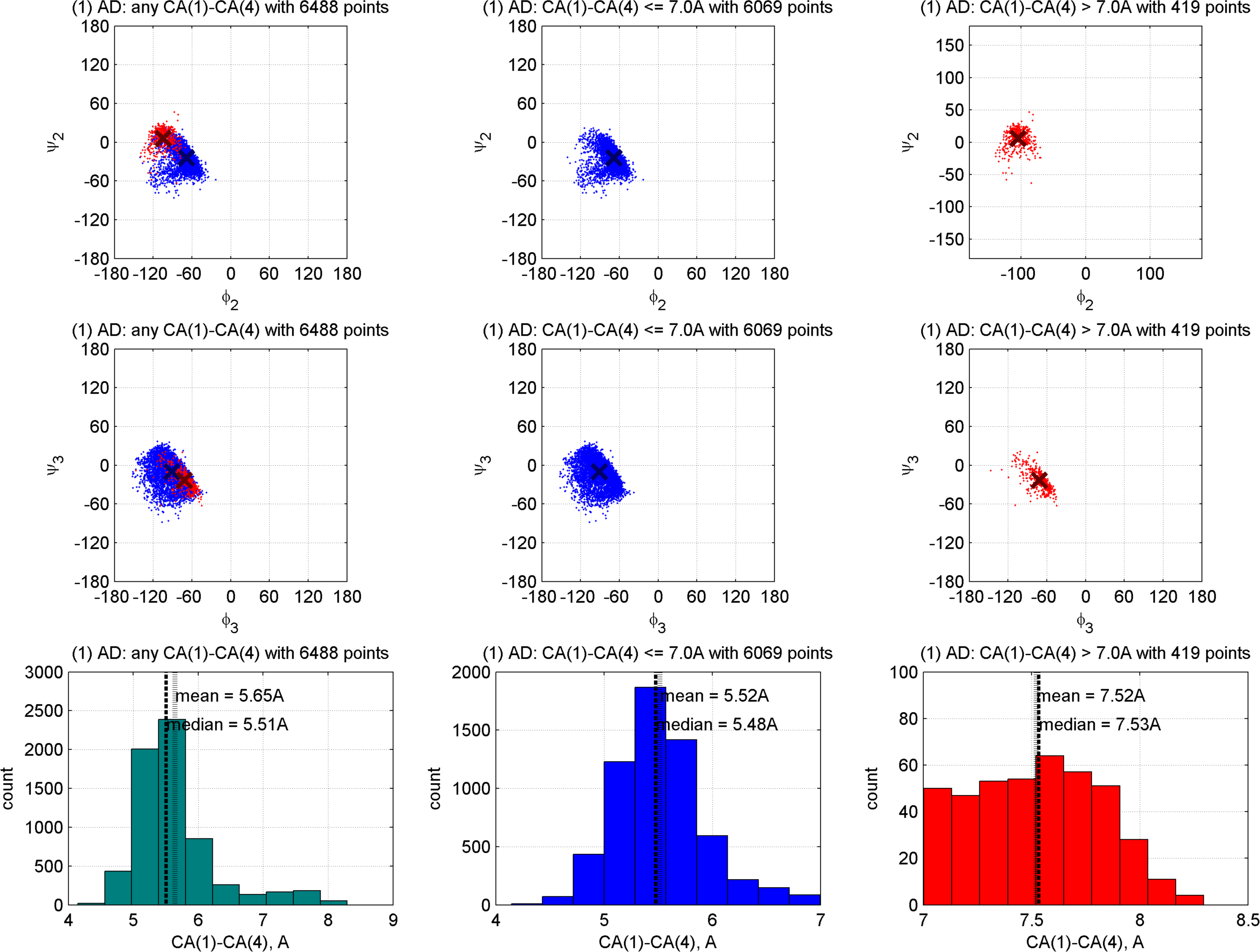
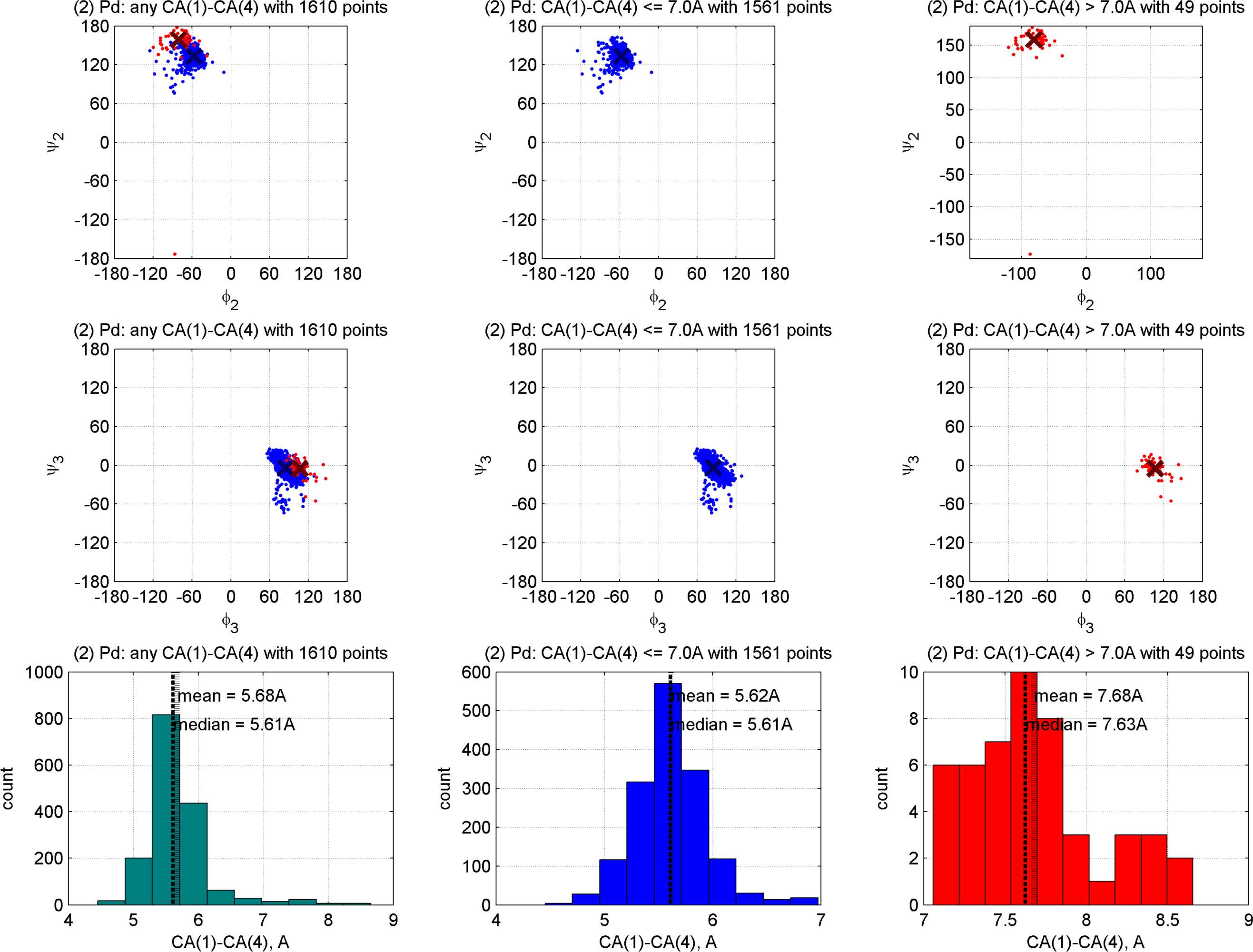
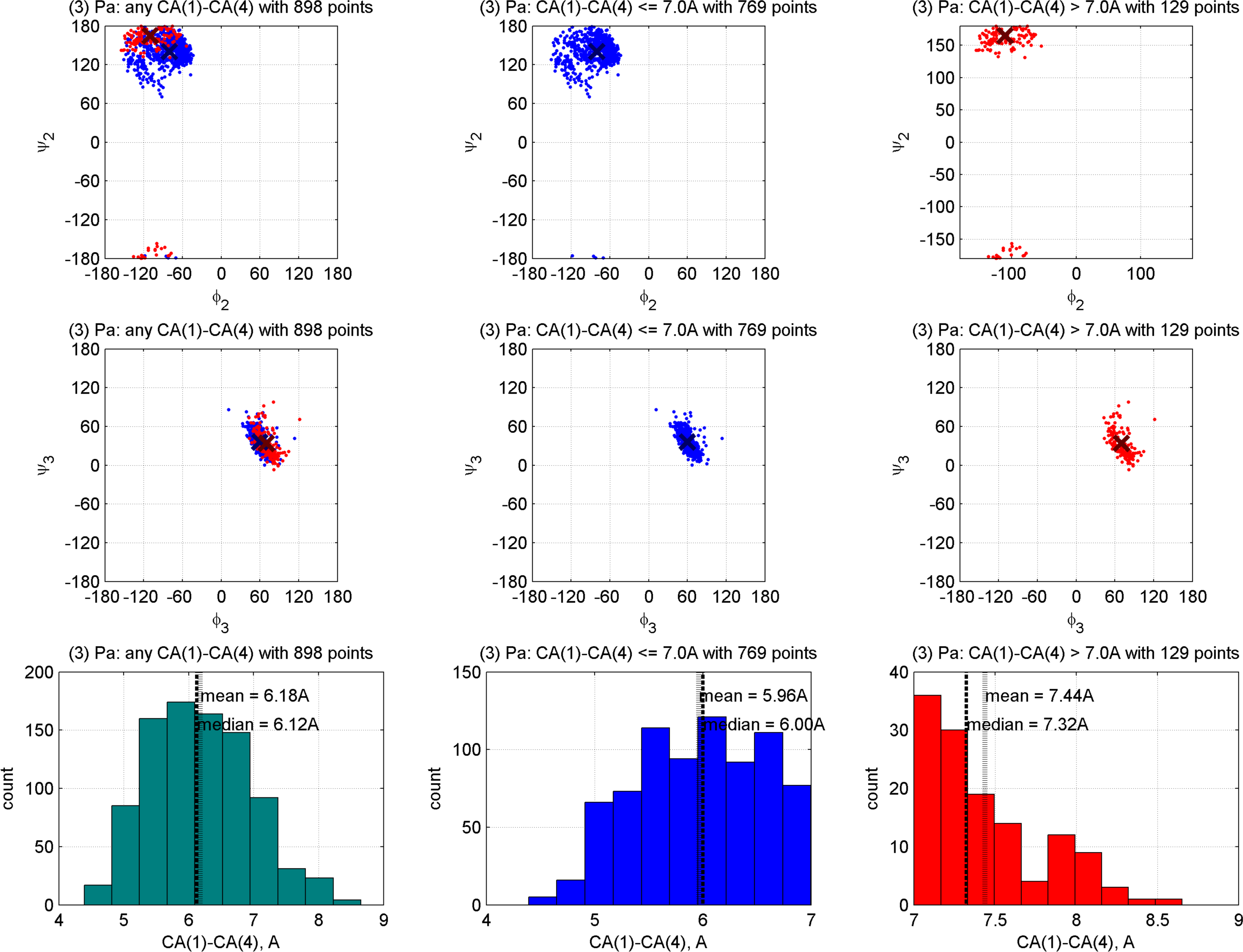
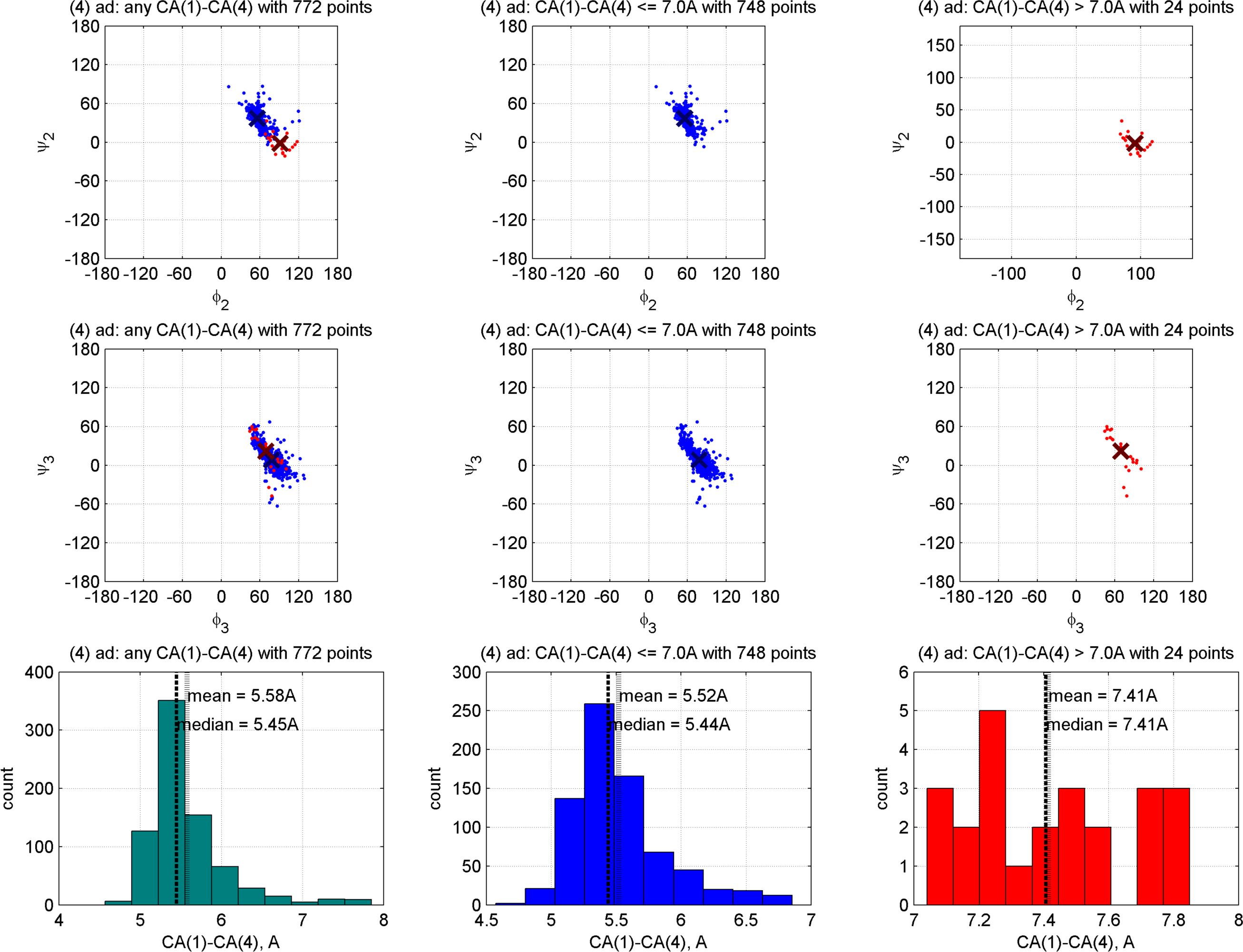
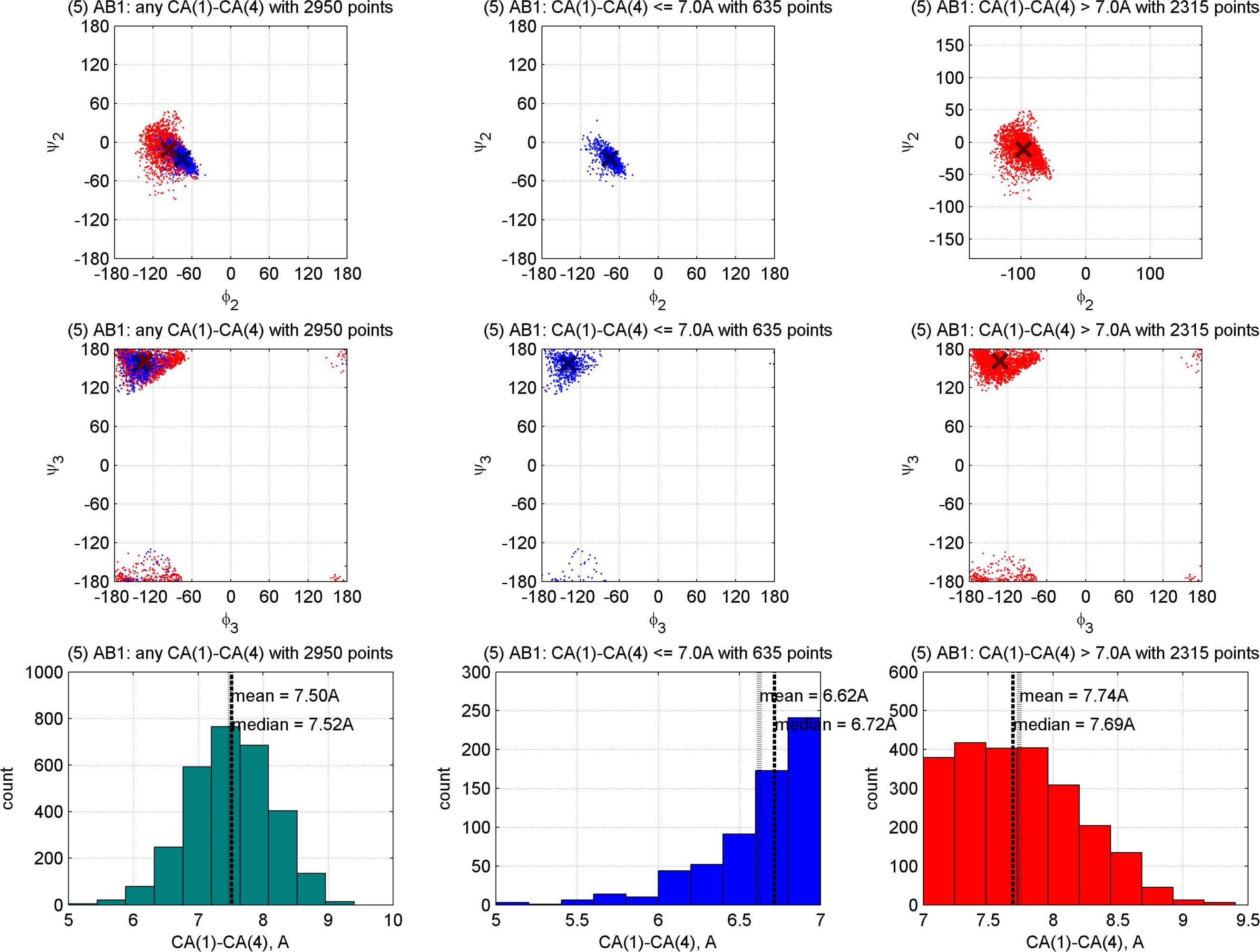
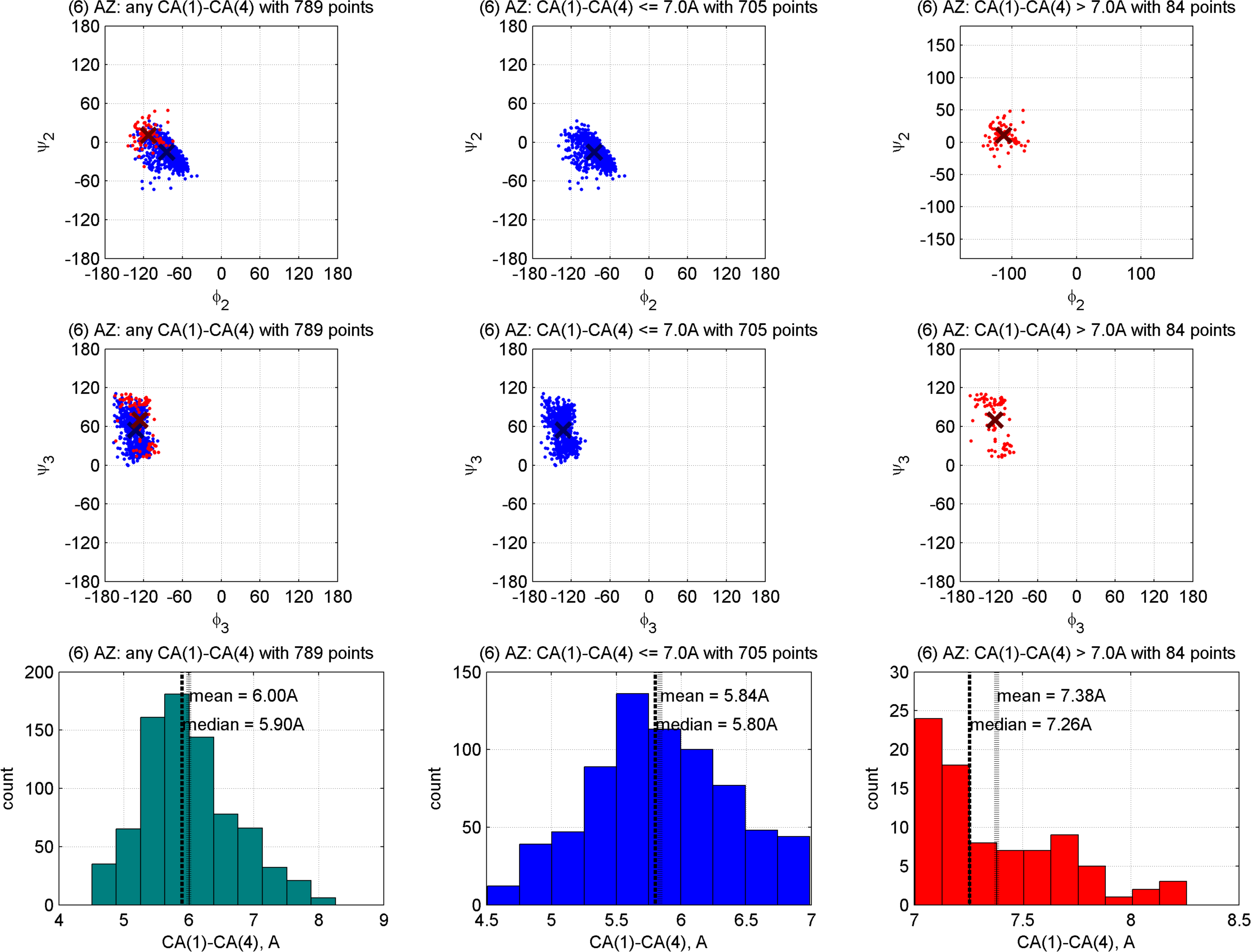
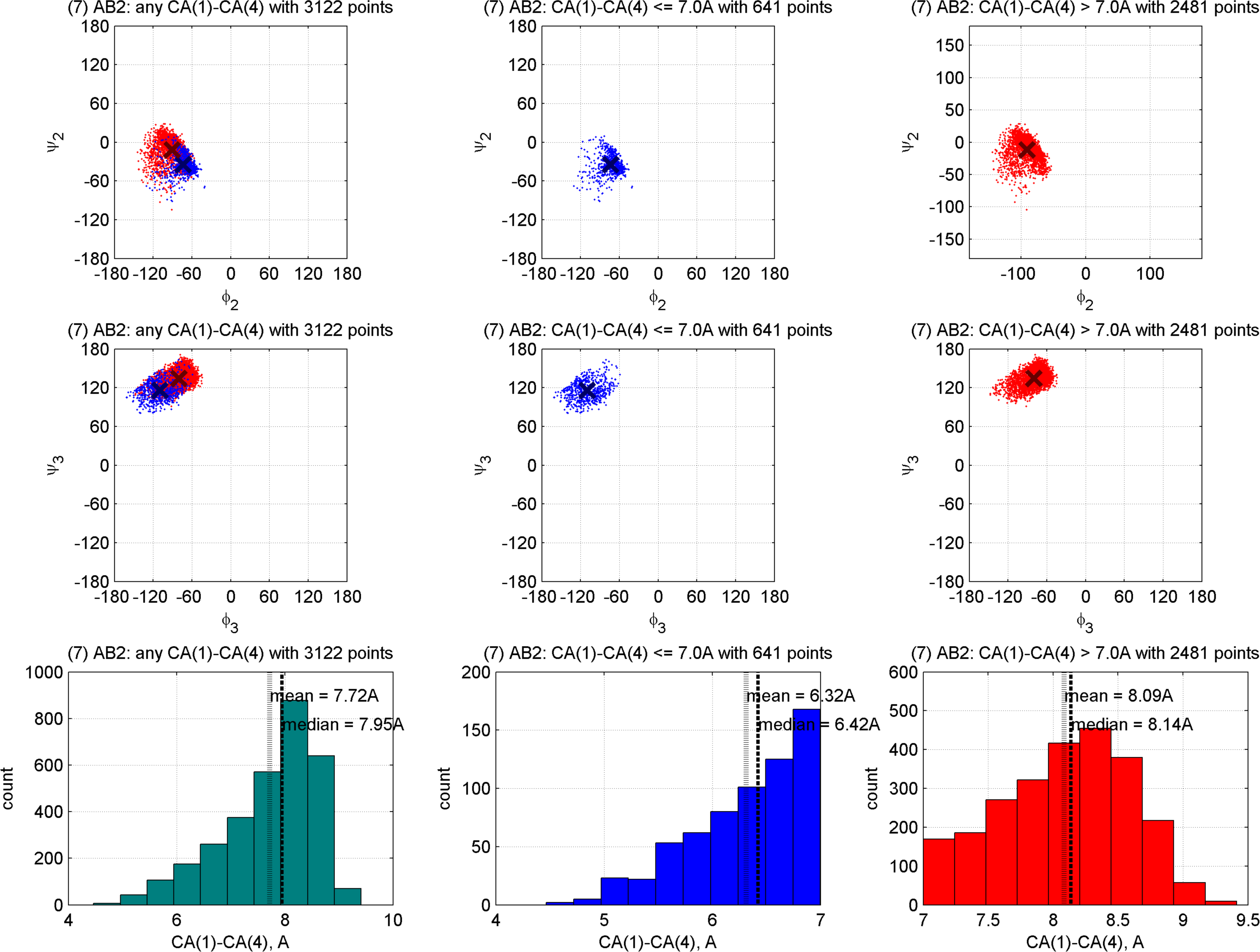
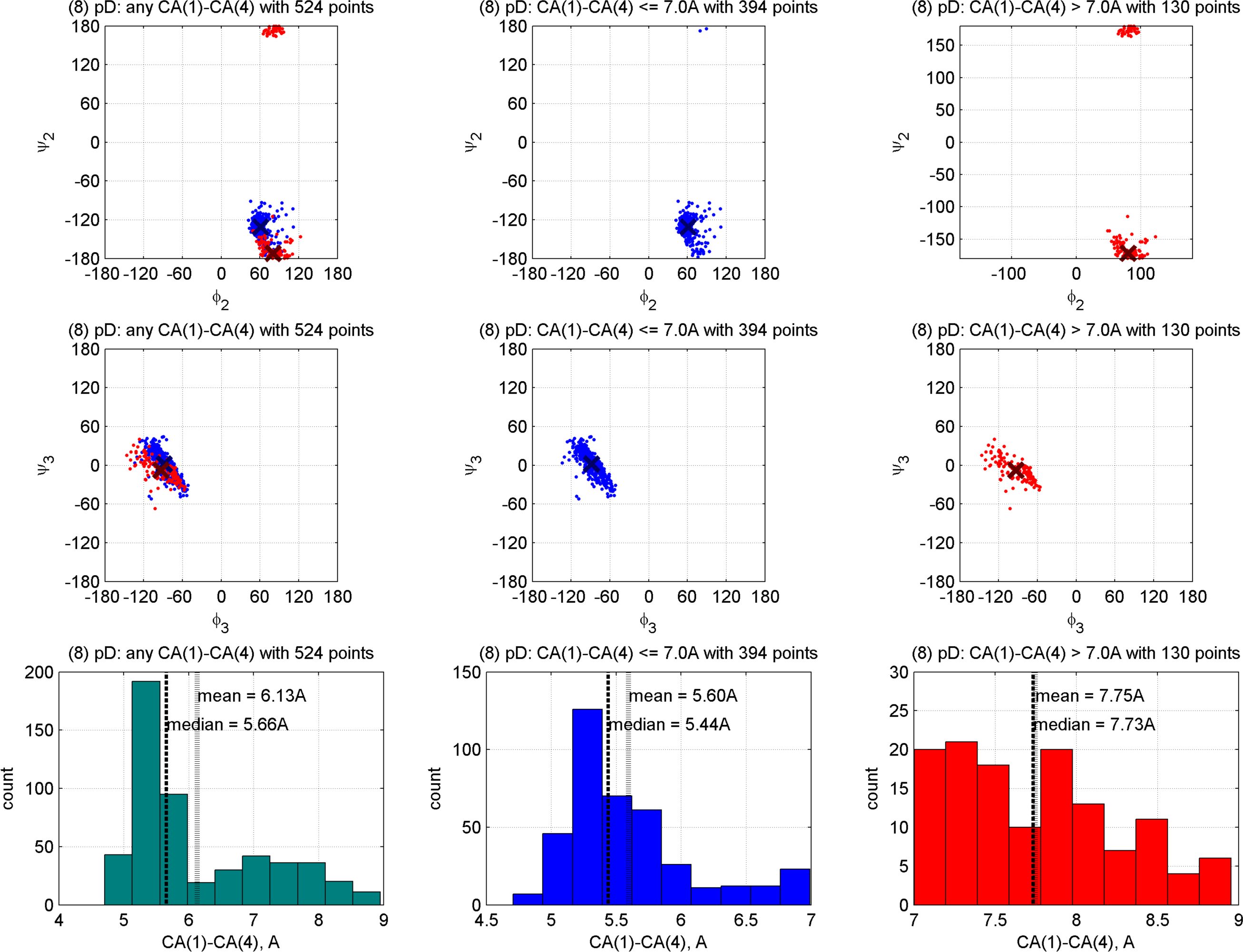
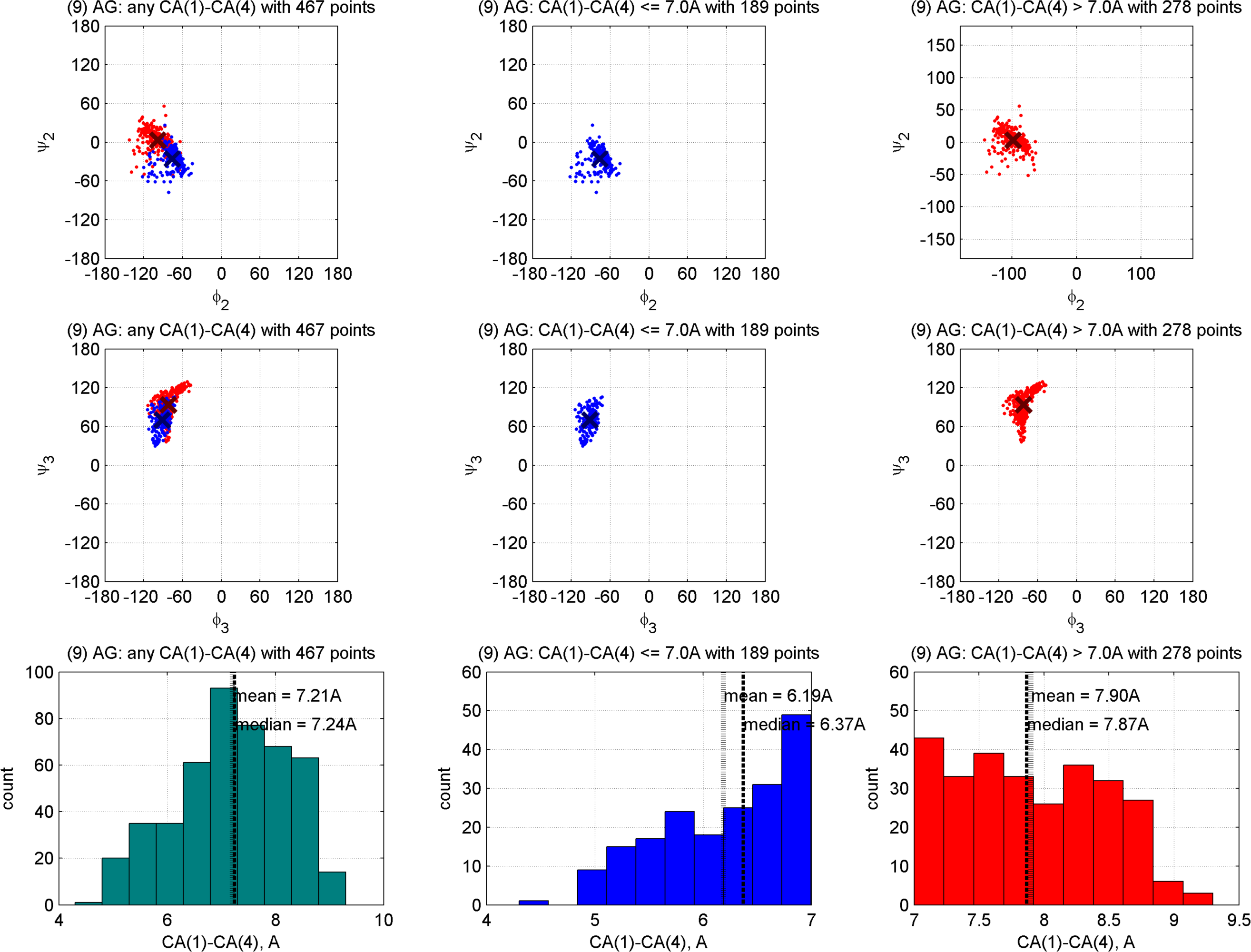
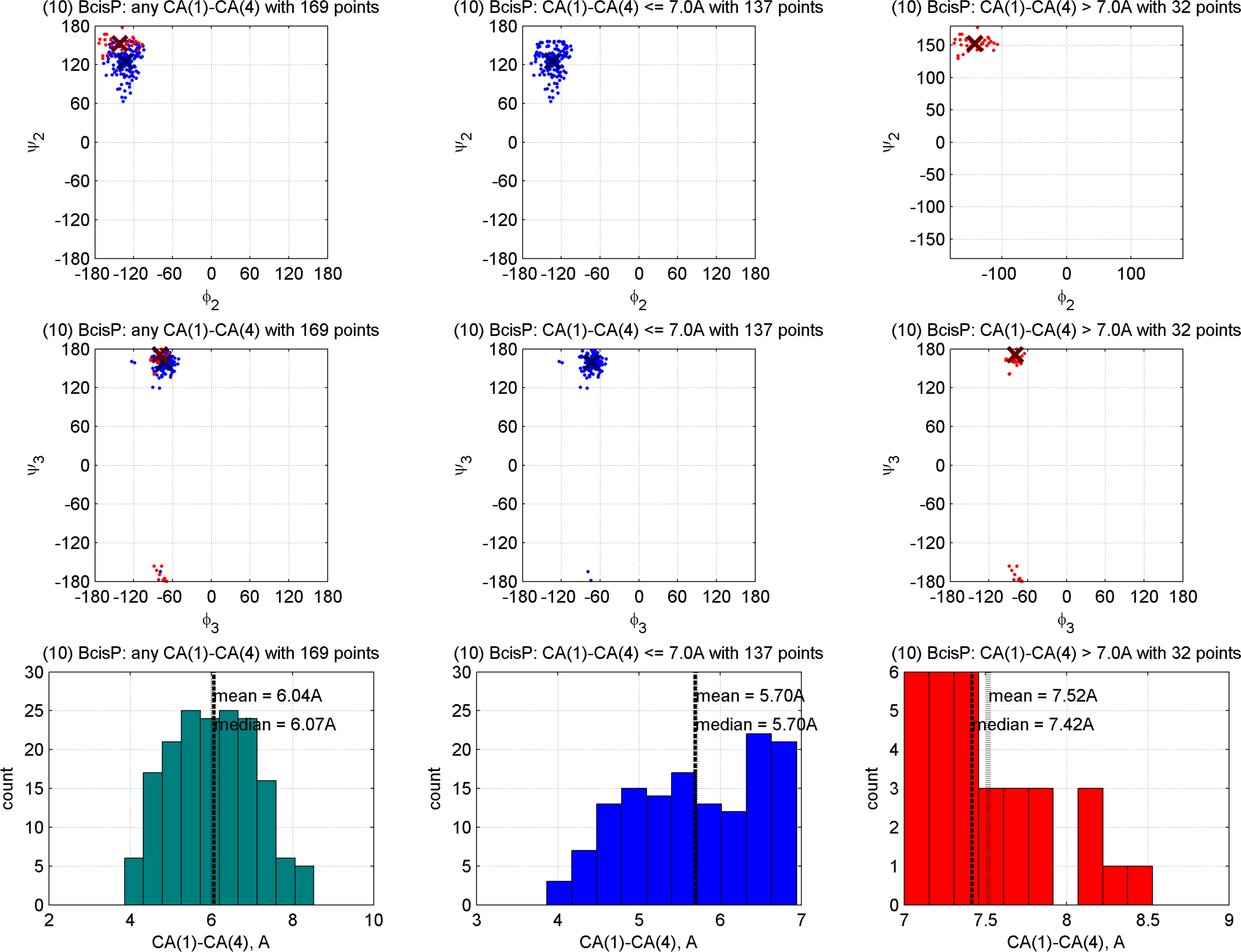
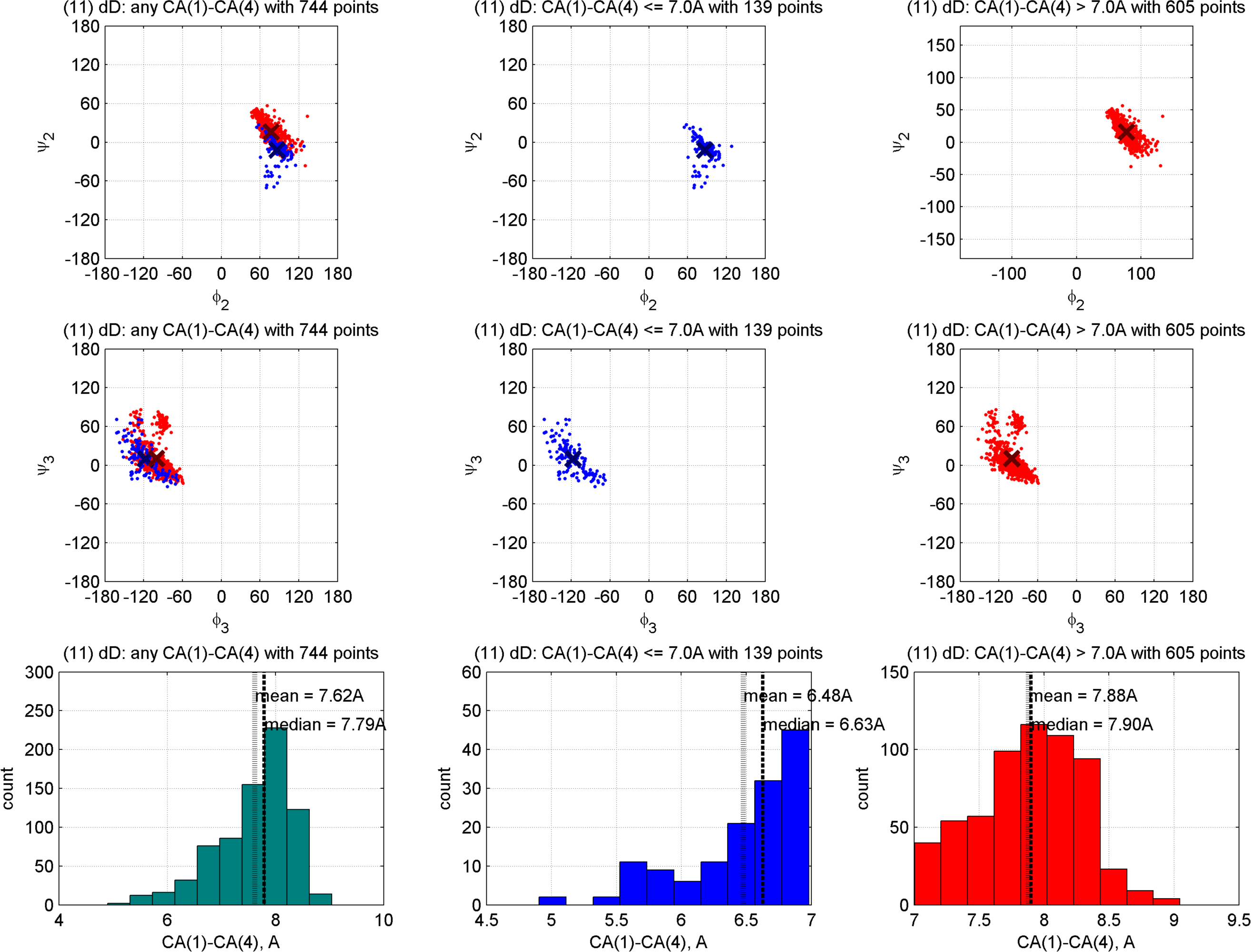
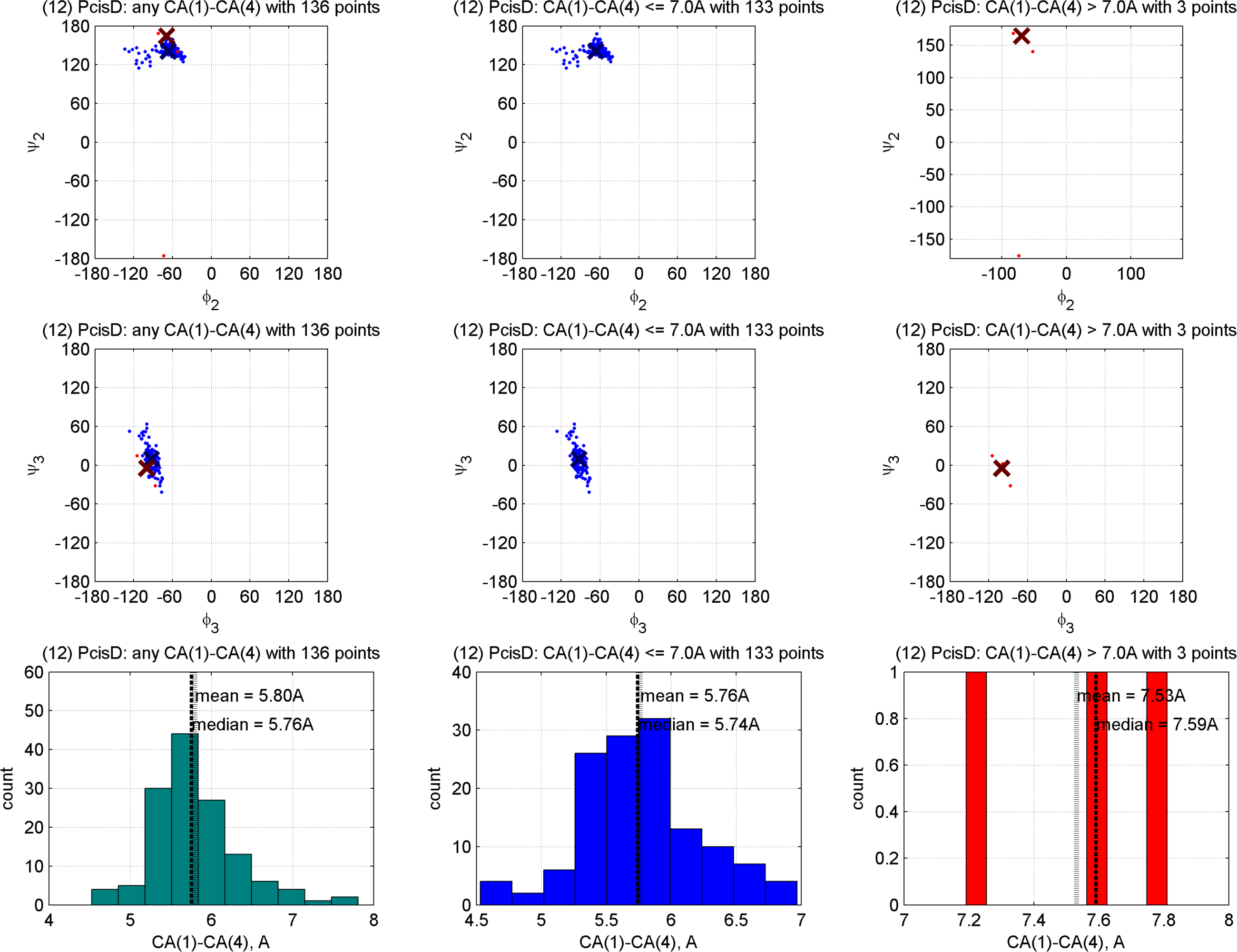
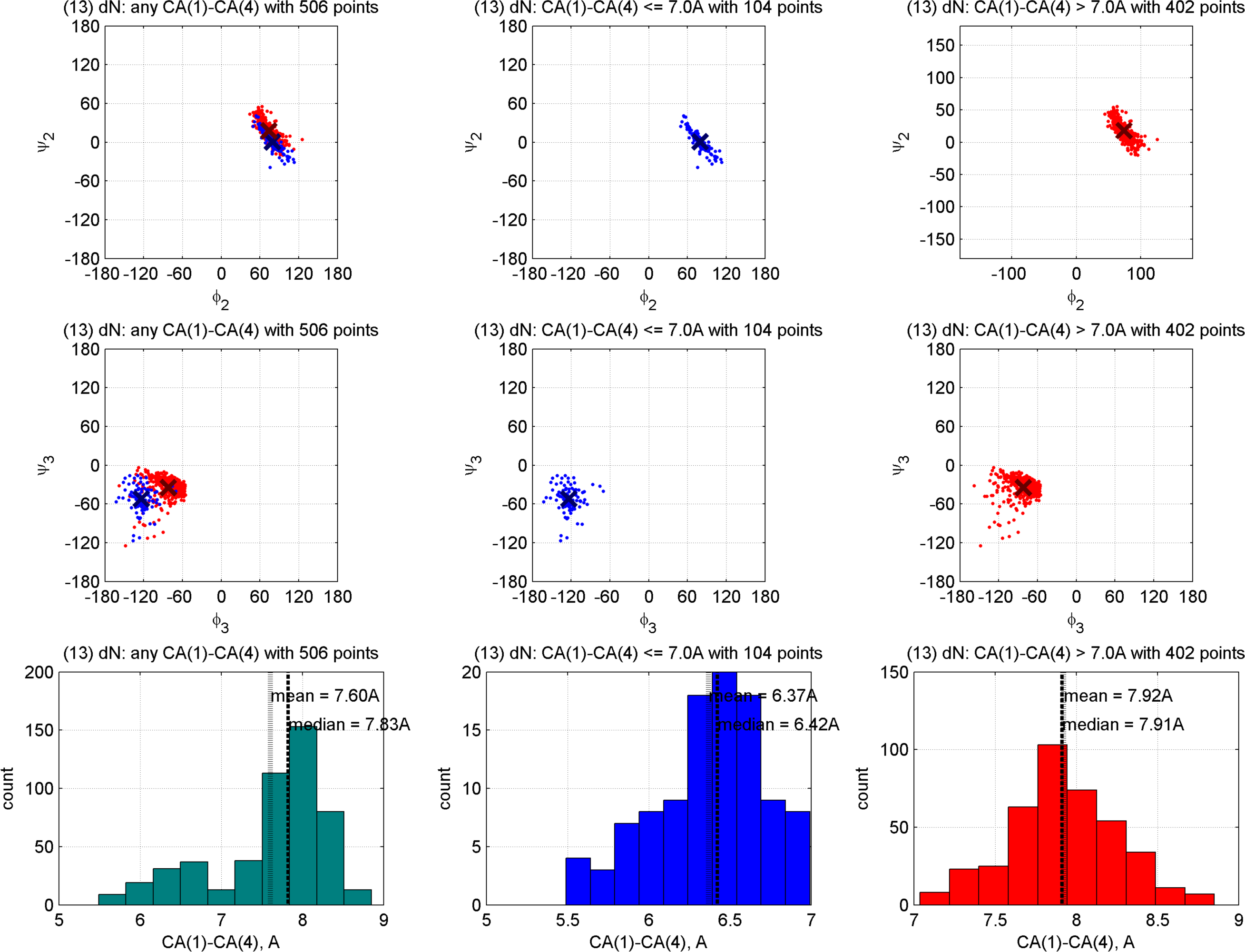
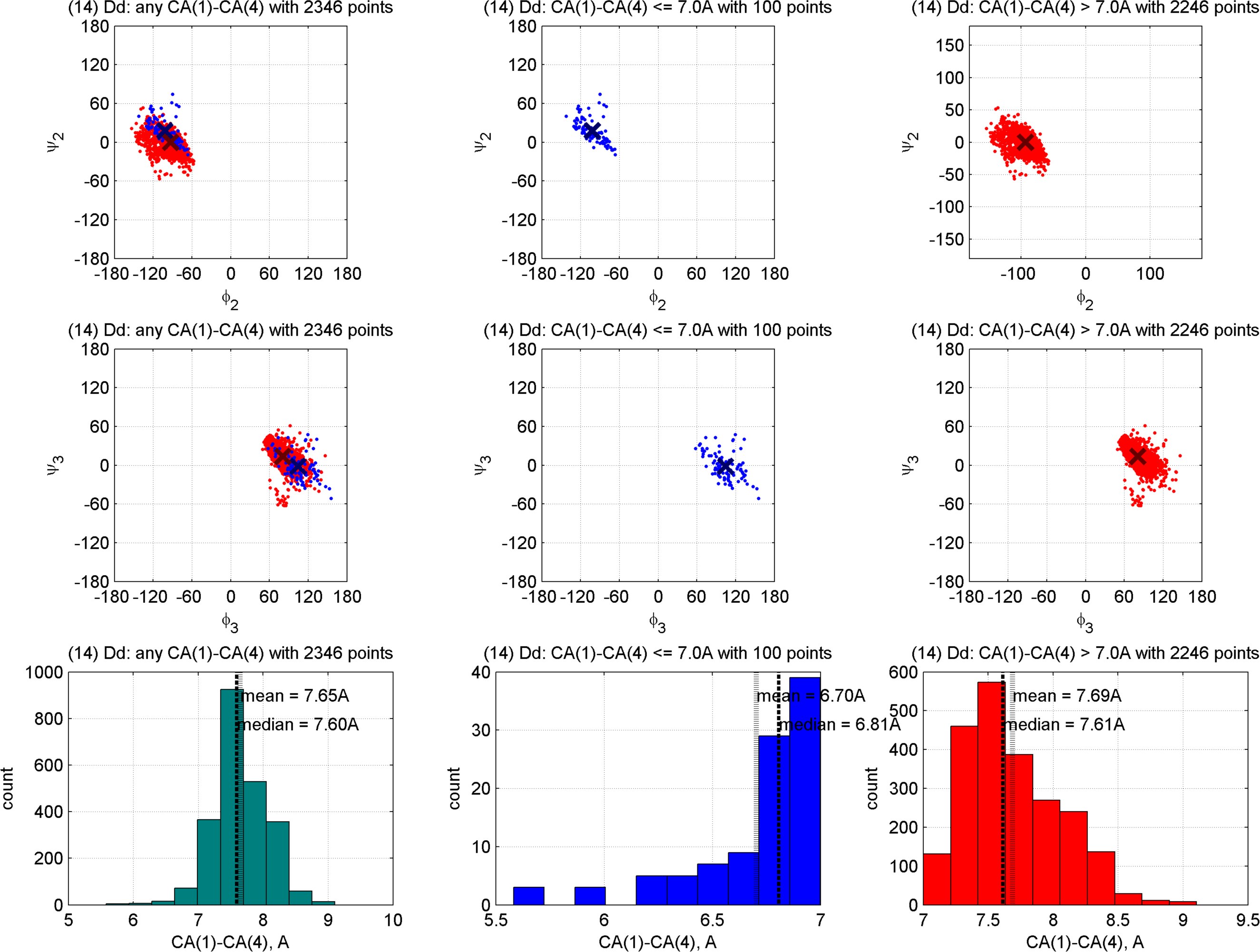
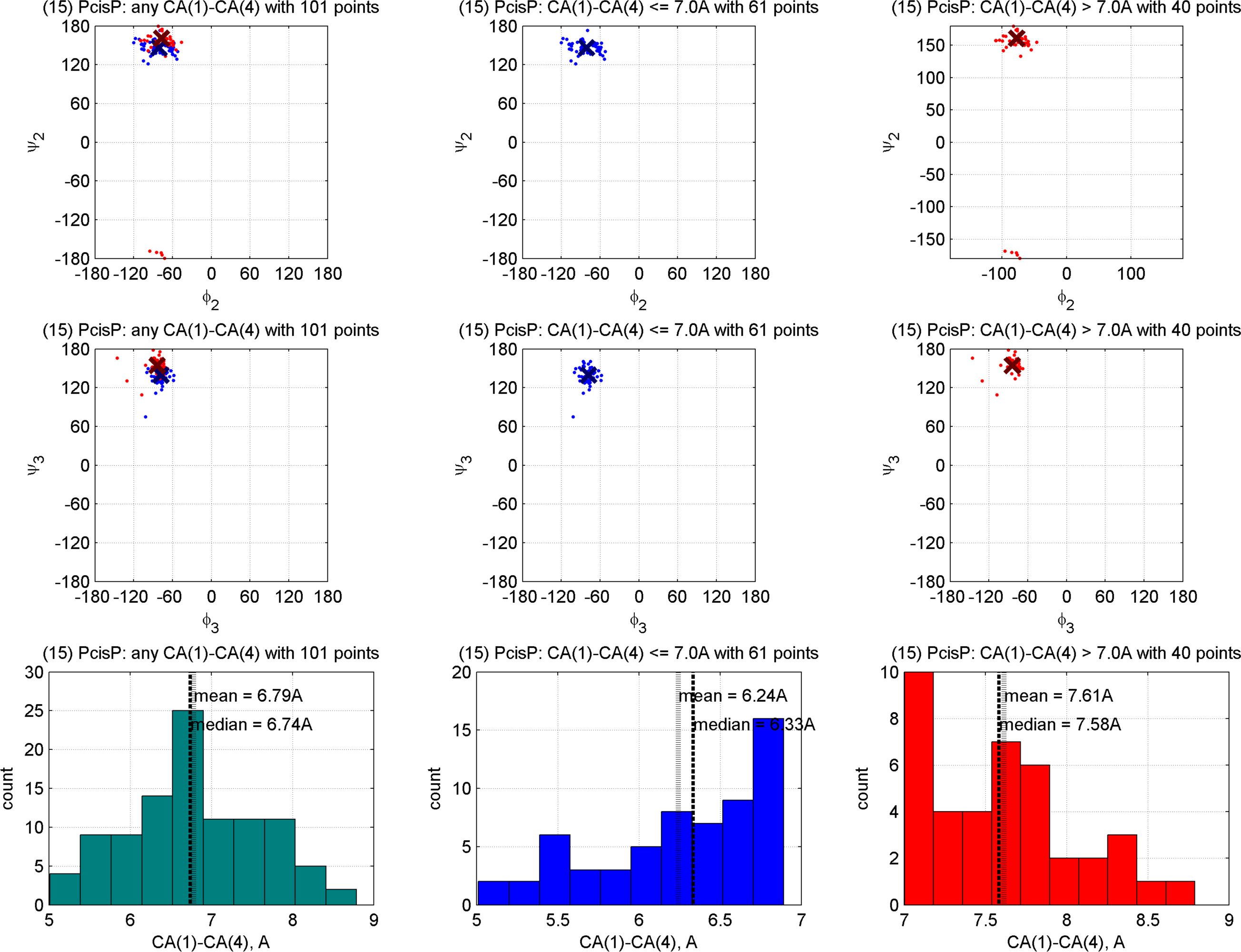
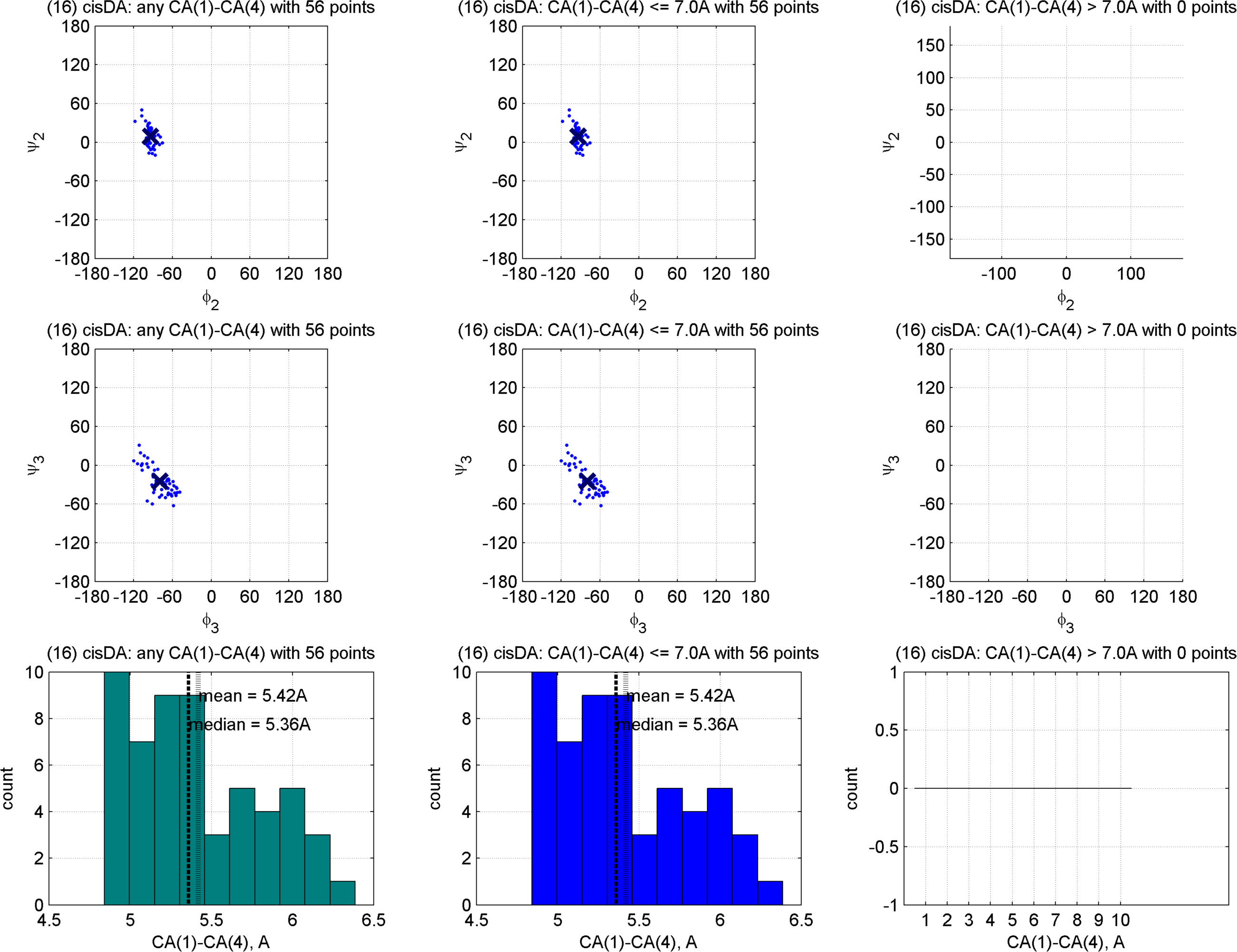
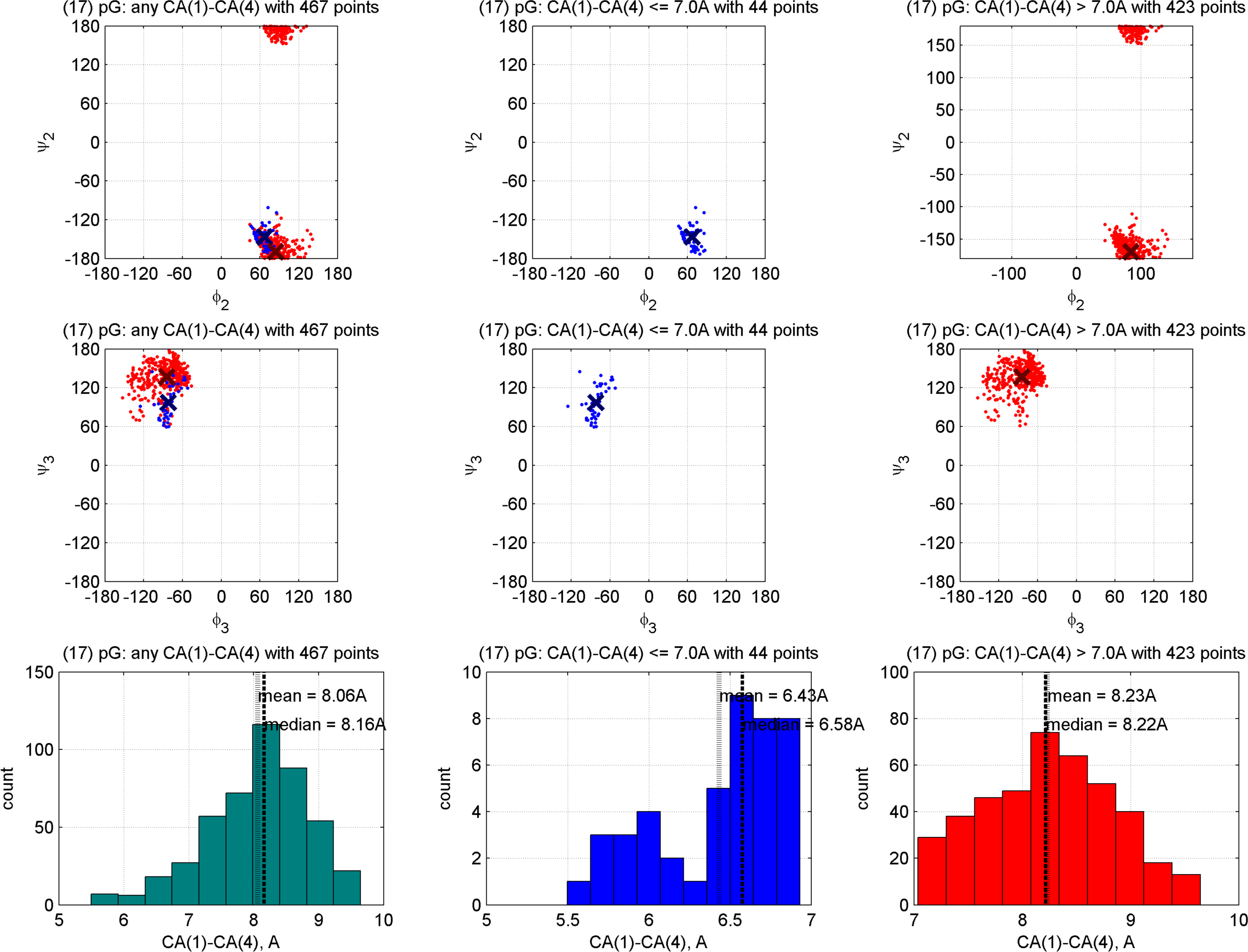
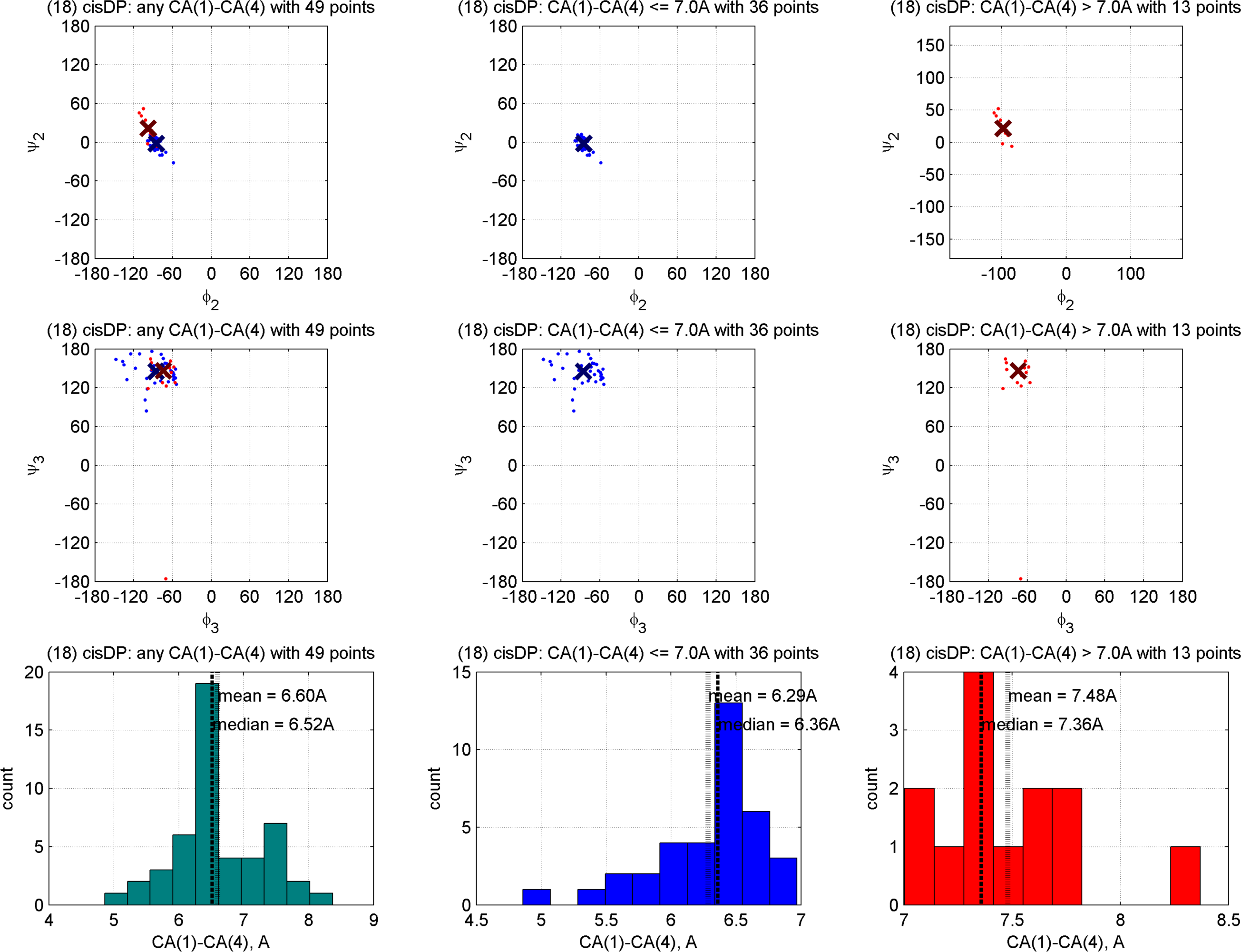
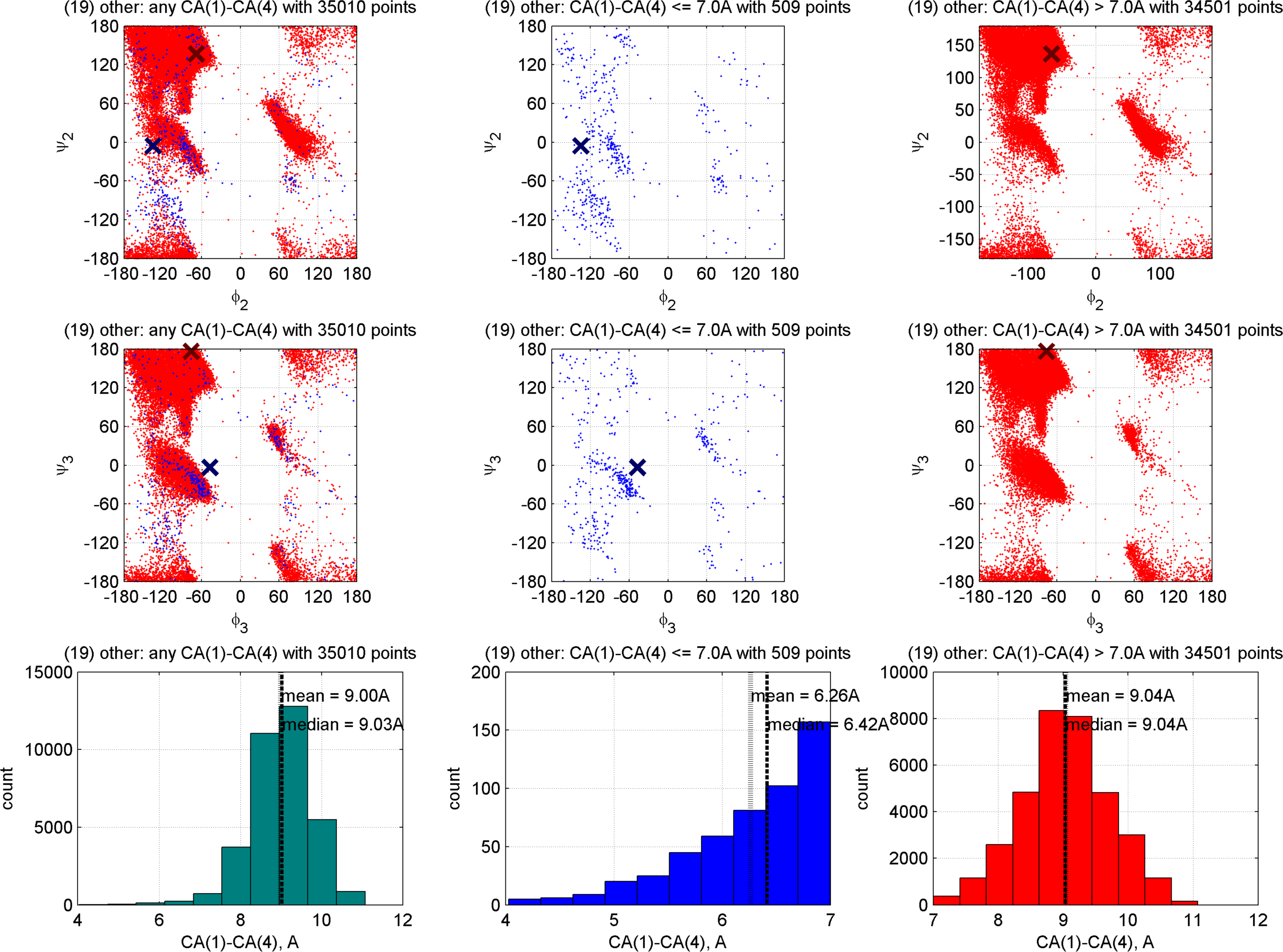

